# Soluble and insoluble dietary fibres differentially affect liver steatosis and gut microbiota in western-diet fed mice

**DOI:** 10.1101/2025.11.26.690531

**Authors:** Céline Marie Pauline Martin, Mathilde Miquel, Valérie Alquier-Bacquie, Arnaud Polizzi, Frédéric Lasserre, Marine Huillet, Clémence Rives, Justine Bruse, Julie Jarassier, Prunelle Perrier, Jelskey Gomes, Claire Naylies, Charlène J. G Dauriat, Nancy Geoffre, Anaëlle Durbec, Adeline Duchampt, Alain Thirault, Xavier Blanc, Elodie Rousseau-Bacquie, Yannick Lippi, Justine Bertrand-Michel, Cécile Canlet, Benoit Chassaing, Anne Fougerat, Laurence Gamet-Payrastre, Amandine Gautier-Stein, Hervé Guillou, Nicolas Loiseau, Sandrine Ellero-Simatos

## Abstract

**Scope:** Metabolic dysfunction-associated steatotic liver disease (MASLD) is the most common chronic hepatic liver disease. An imbalance diet, rich in lipids and sugars and low in fibre, is a key contributing factor. However, there is limited knowledge about how soluble and fermentable dietary fibres, compared to insoluble and non-fermentable fibres, differently affect liver metabolism through their interactions with the intestinal microbiota.

**Methods and results:** Male mice housed at thermoneutrality were fed a Western Diet (WD) supplemented with fermentable inulin or non-fermentable cellulose for 18 weeks. Inulin supplementation mitigated WD-induced obesity, glucose intolerance, dyslipidemia and protected against WD-induced hepatic steatosis compared to cellulose. Hepatic gene expression changes induced by WD were attenuated with inulin. Additionally, inulin preserved gut microbiota composition and metabolism, indicating greater resilience against diet-induced perturbations.

**Conclusion:** These findings suggest that soluble dietary fibres like inulin confer superior metabolic and hepatic benefits over insoluble fibres by modulating the gut microbiota-liver axis, highlighting their potential role in MASLD management.

## Introduction

Metabolic dysfunction-associated steatotic liver disease (MASLD) is the most common chronic hepatic liver disease, with a global prevalence of 30%^[1]^, and is recognized as a major public health issue. MASLD is closely associated with metabolic dysfunctions – including insulin resistance, type 2 diabetes, and obesity – hence the recent name change from non-alcoholic fatty liver disease (NAFLD) to MASLD^[2–4]^. It comprises several pathologies – ranging from steatosis, which is benign and reversible, to the more severe steatohepatitis (MASH), which is characterized by inflammation, hepatocyte damages, and progressive fibrosis, and is a predisposing factor for cirrhosis and hepatocellular carcinoma^[5,6]^. The development and progression of MASLD is largely attributed to an excess calorie intake, mostly in the form of carbohydrates and fat, which enter the liver, resulting in triglyceride accumulation. Although several studies are underway to explore new insights into the prevention and therapy of MASLD, the management of MASLD currently relies on adopting favourable lifestyle and dietary changes^[7]^. Official recommendations include improving diet quality (similar to the Mediterranean dietary pattern), limiting the consumption of ultra-processed food (rich in sugars and saturated fat) and avoiding sugar-sweetened beverages^[8]^. The beneficial effect of the Mediterranean diet on MASLD outcomes is thought to result from a combination of specific nutrients including fruits and vegetables rich in antioxidants, vitamins and polyphenols, olive oil rich in monounsaturated fatty acids and whole grains and nuts, rich in dietary fibres^[9]^.

Dietary fibres are primarily carbohydrates found in the edible part of plant products (i.e., fruits, vegetables, legumes, nuts, seeds and grains) that resist digestion by intestinal and pancreatic enzymes. Numerous controlled trials and meta-analyses demonstrate that dietary fibre supplementation improves metabolic health, including body weight^[10]^, glycaemic control^[11]^ and insulin sensitivity^[12]^ in both healthy and at-risk populations. Similarly, several reviews emphasize that diets rich in whole grains, vegetables, fruits and other fibre sources can improve liver enzymes^[13,14]^ and decrease liver lipid content^[15,16]^. Thus, most evidence in MASLD comes from studies of broader dietary interventions where increased fibre is one component, but trials evaluating specific effects of fibre supplementation are sparse. A recent clinical trial investigated the effects of resistant starch in Chinese MASLD adults and showed that 4 months of resistant starch intervention reduced intra-hepatic lipid content, even after adjusting for weight loss^[17]^. Resistant starch is a type of nondigestible fibre that acts as a prebiotic, being fermented by the microbiota in the large intestine.

Accumulating evidence indeed suggest that MASLD is a disease closely related to gut microbiota via the gut-liver axis^[18]^, and modulating the gut microbiota appears as a promising strategy to improve disease severity^[19]^. Thus, prebiotics, defined as dietary substrates that stimulate the growth of specific taxa, have been extensively studied in humans and preclinical models and shown to reduce insulin resistance, intrahepatic lipids and liver enzymes^[17,20,21]^. Among these prebiotics, dietary fibres are among the most studied and both soluble (e.g., inulin, psyllium, β-glucan) and insoluble (e.g., cellulose, wheat bran) fibres can benefit metabolic health, through different mechanisms. Soluble fibres are fermented by bacterial enzymes within the colon, yielding metabolically active compounds among which short chain fatty acids (SCFA). They are especially effective for improving glycemic control, lowering cholesterol, and reducing body weight and fat^[17,22]^ and these benefits are thought to be due to their viscosity, gel-forming ability, benefic changes in gut microbiota composition and fermentation to SCFA and other metabolites^[23,24]^. Insoluble fibres are generally poorly metabolized by microbial enzymes but still contribute to overall gut and metabolic health^[25]^. They are also consistently linked to reduced risk of type 2 diabetes and improved insulin resistance in long-term cohort studies^[26,27]^ and are thought to act mainly by increasing stool bulk and reducing intestinal transit time, with additional anti-inflammatory and metabolic effects^[28]^. However, there is less evidence regarding whether the two types of fibres differentially influence liver metabolism and MASLD progression, and most studies have been conducted using high-fat-diet (HFD) models of metabolic perturbations.

Compared to HFD, mainly enriched in lipids, we previously demonstrated that a Western-Diet (WD), enriched in lipids, cholesterol and sucrose, induced significantly more steatosis, liver inflammation and fibrosis in mice^[29]^, thus recapitulating MASH features. A recent unbiased ranking of murine dietary models based on their proximity to human MASLD confirmed that WD aligned closely with human MASH^[30]^, and WD are increasingly recognized as some of the best models in recapitulating human MASLD^[31]^. Furthermore, we have previously shown that, compared to standard temperature housing, housing mice at thermoneutral conditions (30°C) induced more severe steatosis and steatohepatitis in mice fed a WD. We have also shown that thermoneutral housing enabled a more homogenous response to diet compared to standard temperature housing^[32]^. In the present study, we therefore used thermoneutral housing in combination with a WD, a relevant nutritional model of MASLD, to investigate the effects of dietary supplementation with two types of dietary fibres: cellulose (not fermentable by gut microbiota) and inulin (fermentable by gut microbiota) on MASLD development.

## Experimental section

### Animals

All experiments were carried out in accordance with the European Guidelines for the Care and Use of Animals for Research Purposes and approved by an independent ethics committee (CEEA-86 Toxcométhique) under the authorization number 16136-2018062810452911 v5. Animals were treated humanely with due consideration to the alleviation of distress and discomfort. A total of 48 C57BL/6J male mice (8-week-old) were purchased from Charles Rivers Laboratories (L’Arbresle, France). All mice were housed on a 12 h light (ZT0-ZT12)/12 h dark (ZT12-ZT24) cycle in a ventilated cabinet at the thermoneutral temperature (TN = 28-30°C) throughout the experiment. Mice were allowed two weeks of acclimatization with free access to water and food (chow diet supplemented with 9% of Cellulose or Inulin) from SAAJ INRAe (Jouy, France, see also Table S1). Then, the mice were randomly divided into four groups of 12 mice. Half of the mice were fed a chow diet (CD) and the other half a Western diet (WD). CD and WD were supplemented with 9% of dietary fibre which was either pure cellulose (9%, thereafter named “cellulose diet”) or a mixture of cellulose (2.25%) and inulin (6.75%, thereafter named “inulin diet”) (SAAJ INRAe, Jouy, see also Table S1). Dietary intervention lasted 18 weeks and each experimental group was composed of 3 independent cages of 4 mice. Individual mouse body weight, and food and water intakes per cage were measured weekly. Fresh feces were collected after 13 weeks of diet for bile acids and NMR metabolomics and 15 weeks of diet for 16S sequencing in the morning (ZT0).

### Oral glucose tolerance test (OGTT) and plasma insulin measurement

All experiments were performed using conscious mice. After 15 weeks of feeding, mice were fasted for 6h before receiving an oral glucose load (2 g/kg body weight). Blood glucose was measured at the tail vein using an Accu-Check® Performa glucometer (Roche Diabetes Care France, Mylan, France) at 30 min before and 0, 15, 30, 60, 90 and 120 min after the glucose load. At 30 min before and 15 min after glucose injection, 20 µl blood from the tip of the tail vein was sampled for measurement of plasma insulin concentration using HTRF serum kits (Cisbio, Codolet, France).

### Blood and tissue sampling

Prior to sacrifice, mice were anaesthetised with isoflurane and xylazine (2%, 2mg/kg) then blood from the vena cava and the portal vein was collected into lithium heparin-coated tubes (BD Microtainer®, BD, Dutscher, Brumath, France). Then, mice were euthanised by cervical dislocation. Plasma was isolated by centrifugation (1500 × g, 15 min, 4°C) and stored at -80°C until biochemical analysis. Tissue samples (liver, subcutaneous and perigonadic white adipose tissue (WAT), caecal content) were collected, weighed and snap-frozen in liquid nitrogen and stored at −80°C.

### Plasma biochemical analysis

Plasma samples were assayed for alanine transaminase (ALT), alkaline phosphatase (ALP), free fatty acids (FFAs), triglycerides (TGs), total cholesterol, low-density lipoprotein (LDL-Cholesterol) and high-density lipoprotein (HDL-Cholesterol). All biochemical analyses were performed with a ABX Pentra 400 biochemical analyzer (Anexplo facility, Toulouse, France).

### Liver neutral lipid analysis

Hepatic lipids were extracted and liver cholesterol, cholesterol esters and triglycerides were measured as previously described^[32]^.

### Histology

Paraformaldehyde-fixed, paraffin-embedded liver tissue was sliced into 5 µm sections and stained with hematoxylin and eosin (H&E) or with Sirius red or used for immuno-histochemistry. The staining was visualized with a light microscopy equipped with a Nikon 90i. All liver sections were analysed blindly. Liver steatosis was evaluated according to Kleiner^[33]^. Sirius Red and anti α-smooth muscle actin (α-SMA) stainings were performed as previously^[32]^.

### Gene expression

Microarray experiments were conducted on n = 7 mice per group (randomly chosen). Gene expression profiles were performed at the GeT-TRiX facility (GénoToul, Génopole Toulouse Midi-Pyrénées, France) using Sureprint G3 Mouse GE v2 microarrays (8 x 60K, design 074809, Agilent Technologies) following the manufacturer’s instructions and as described in^[32]^. Microarray data are available in the ArrayExpress database (http://www.ebi.ac.uk/arrayexpress) under accession number E-MTAB-16279.

Microarray data analyses were performed as previsouly^[32]^. Briefly, probes with fold-change > 1.5 and FDR ≤0.05 were considered to be differentially expressed between conditions.

### Fecal microbiota composition analysis by 16S rRNA gene sequencing using Illumina technology

16S ribosomal RNA (rRNA) gene amplification and sequencing were performed as described previously^[34]^ using an Illumina MiSeq sequencer (paired-end reads, 2x250 bp) at the Genom’IC facility at Cochin Institut, Paris, France. 16S rRNA sequences were analysed with QIIME2 – version 2024.10^[35]^. Sequences were demultiplexed and quality-filtered with the Dada2 method. QIIME2 default parameters were used to detect and correct Illumina amplicon sequence data, and a table of Qiime 2 artifacts was generated. Next, a tree was generated with the align-to-tree-mafft-fasttree command, for phylogenetic diversity analyses. Alpha and beta diversity analyses were computed with the core-metrics-phylogenetic command. Principal coordinates analysis (PCoA) plots were constructed to assess the variation between the experimental groups (β-diversity). α-diversity was computed with the Shannon, Simpson, Observed species, Chao1 and Inverse Simpson indexes. To analyse the taxonomy, features were assigned to operational taxonomic units (OTUs), according to a 99% threshold of pairwise identity to the Silva 138 reference database. Unprocessed sequencing data are deposited in the European Nucleotide Archive under accession number XXXX. Statistical analyses for the differential abundances of phylum and genus were performed using the methodology ANCOM-BC^[36]^.

### Proton nuclear magnetic resonance (^1^H-NMR)-based metabolomics

Fecal samples collected at the end of the experiment were prepared and analyzed using ^1^H-NMR-based metabolomics. All spectra were obtained on a Bruker DRX-600-Avance NMR spectrometer (Bruker) on the AXIOM metabolomics platform (MetaToul). Details on experimental procedures, data pre-treatment and statistical analysis were described previously^[37]^. For illustration purposes, the area under the curve (AUC) of several signals of interest was integrated and significance tested with one-way ANOVA as described below.

### Bile acid extraction and analysis

Fecal samples collected at the end of the experiment were used. 30 mg of feces were crushed into 1 ml of sodium hydroxyde (0.2M). 40 µL of internal standard (23-Nordeoxycholic acid) were added to 3 mg of feces into a 1.5 ml glass tube. Samples were incubated 1 hour at 60°C and then cooled 5 min at room temperature. 2 ml of H_2_O were added and sample were centrifuged 15min at 2500 rpm. 1.5ml of samples were then submitted to solid phase extraction (SPE) using OASIS HLB 96-well plate (30 mg/well, Waters, Milford, MA, USA) conditioned with MeOH (1 mL) and equilibrated with H_2_O (2 mL). After sample application, extraction plate was washed with H_2_O (2 mL). the plate was dried under nitrogen, and bile acids were eluted with 2mL of MeOH. Prior to LC-MS/MS analysis, samples were evaporated under nitrogen and reconstituted in 20 µL with MeOH. Briefly, bile acids were separated on a BEH C18 column (2.1 mm, 150 mm, 1.8 µm) (Waters, Milford, MA, USA) using high performance liquid chromatography system (Acquity Class I; Waters, Milford, MA, USA) coupled to a ESI-High resolution mass spectrometer (Orbitrap, Thermo Fisher Scientific, Waltham, MA, USA). Data were acquired in full scan mode with optimized conditions. Peak detection, integration and absolute quantitative analysis were done using TraceFinder software (Thermo Fisher Scientific, Waltham, MA, USA) based on calibration lines built with commercially available bile acids standards (Cayman Chemicals, Ann Arbor, MI, USA).

### Other statistical analyses

All data are presented as means ± standard error of the mean (SEM). Statistical analysis on biochemical and qPCR data were performed using GraphPad Prism version 9 for Windows (GraphPad Software, San Diego, CA). One-way ANOVA was performed followed by appropriate post-hoc tests (Sidak’s multiple comparisons test) when ANOVA was found to be significant (p<0.05). For histological scores, non-parametric test (Kruskall-Wallis) were used. Significance was indicated by: * or # for p < 0.05, ** or ## for p < 0.01, *** or ### for p < 0.001, **** or #### for p < 0.0001.

### Correlation analysis

Integrative analysis of phenotypical data with microbiota taxa at the genus level or with fecal bile acids was performed in R (version 4.4.2) using spearman correlation and regularized canonical correlations analysis (rCCA) with the mixOmics package (version 6.30.0)^[38]^. Prior to integrative analysis with other datasets, the microbiota dataset was filtered to keep only genera with a minimal total count of 0.01% and a minimal prevalence of 25% in at least one experimental group. It was then processed according to the multivariate statistical mixMC framework, which included data processing using Total Sum Scaling (TSS) normalization and log-ratio transformation. The fecal bile acid dataset was also processed to keep only bile acids with a minimal prevalence of 25% in at least one experimental group, and missing data were imputed with a small random value multiplied by the limit of detection for that metabolite.

## Results

### Inulin alleviated WD-induced obesity, glucose intolerance and dyslipidemia in mice

To investigate the effects of insoluble vs. soluble fibres on diet-induced MASLD, male mice were fed a chow (CD) or western-diet (WD) supplemented with dietary fibres which were either pure cellulose (9%, insoluble, thereafter named “CD-cellulose” or “WD-cellulose”) or a mix of cellulose (2.25%) + inulin (6.75%, soluble, thereafter named “CD-inulin” or “WD-inulin”) for 18 weeks at thermoneutral (TN) housing (30°C) (Figure 1A). After 4 weeks, WD-fed mice were significantly heavier than CD-fed mice regardless of the type of dietary fibres (Figure 1B). At the end of the experiment, WD-fed mice supplemented with inulin developed significantly less obesity than those supplemented with cellulose, as confirmed by body weight gain and relative white adipose tissue weights (Figure 1C-E), despite significantly higher total food intake in WD-inulin compared to WD-cellulose-fed mice (Figure S1). Similarly, all WD-fed mice developed glucose intolerance, but significantly less under WD-inulin than under WD-cellulose (Figure 1F). Nevertheless, WD-feeding induced fasted hyperglycaemia regardless of the type of dietary fibres, while significant fasted hyperinsulinemia was observed only in WD-cellulose compared to CD-cellulose mice (Figure 1H). A similar trend was found in WD-inulin vs. CD-inulin, although not statistically significant. Regardless of type of dietary fibres, WD thus significantly increased the HOMA-IR (Homeostasis Model Assessment of Insulin Resistance) index (Figure 1I) and significantly decreased the QUICKI (quantitative insulin sensitivity check index) (Figure 1J). Plasma biochemical analyses revealed that the elevated levels of HDL-, LDL-, total cholesterol and triglycerides observed in mice fed a WD compared to mice fed a CD were significantly lower in WD-inulin mice compared to WD-cellulose mice (Figure 1K-N).

**Figure 1:**
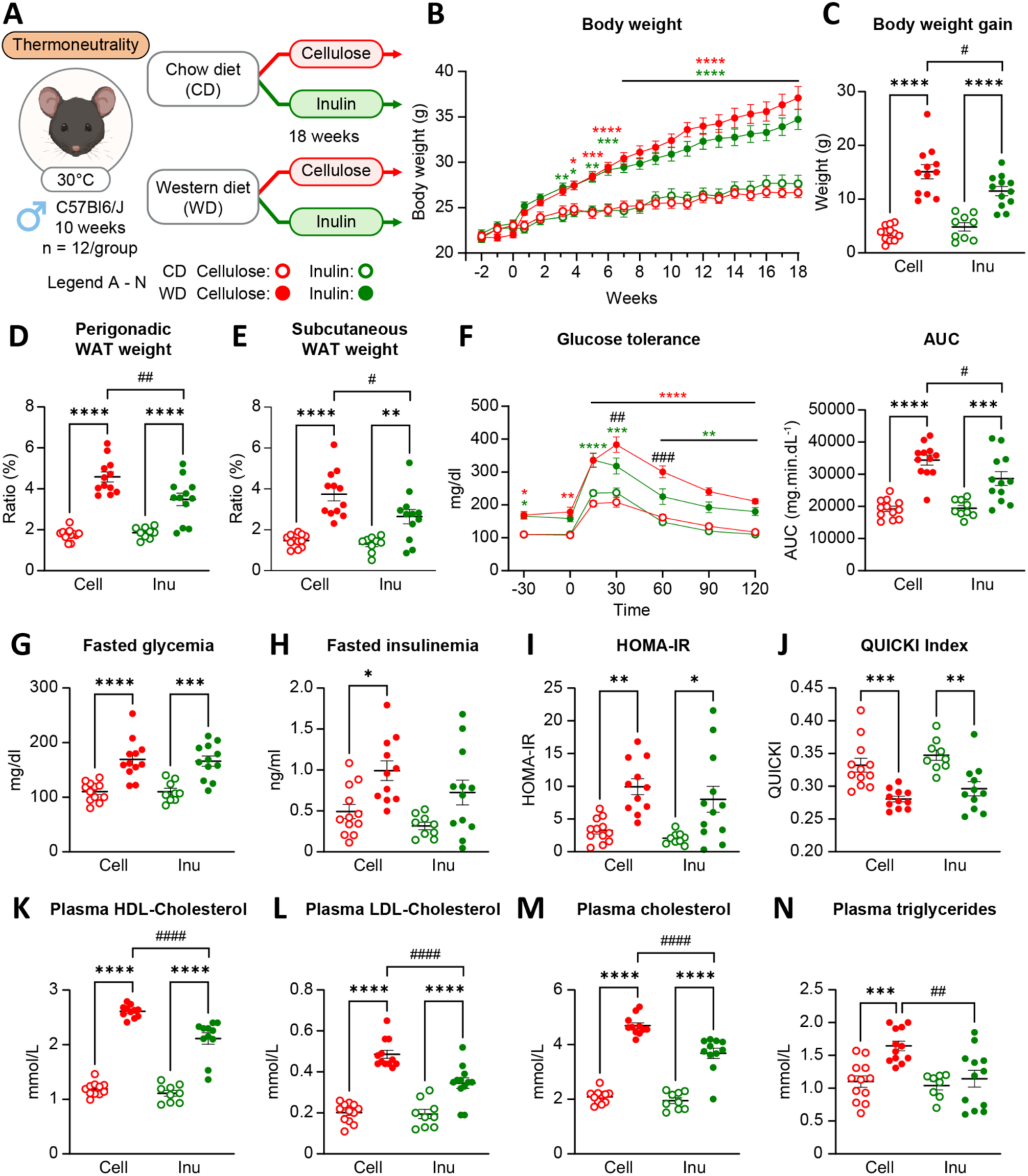
Impact of fibre-enriched WD on obesity, glucose intolerance and dyslipidemia in mice. (A) C57Bl/6 male mice were housed at TN (30°C) and fed a CD or a WD enriched with 9% cellulose or inulin for 18 weeks. (B) Body weight. (C) Body weight gain. (D, E) Ratio of (D) perigonadal and (E) subcutaneous white adipose tissue (WAT) to body weight. (F) OGTT. (G) Fasted glycemia. (H) Fasted insulinemia. (I) HOMA-IR. (J) QUICKI. (K, L, M, N) HDL-cholesterol (K), LDL-cholesterol (L) plasma free cholesterol (M) and triglycerides (N). Data are presented as the mean ± SEM for n = 11-12/group. * diet effect or # fibre effect. Cell = cellulose, Inu = inulin.

Taken together, these results show that the enrichment of WD with inulin alleviated WD-induced obesity which was associated with improved glucose homeostasis and plasma lipid profiles.

### Inulin protected against WD-induced hepatic steatosis

We next investigated the hepatic phenotype. We previously published that male mice housed at thermoneutral housing and fed with a WD containing 5% cellulose developed hepatic steatosis, inflammation and fibrosis after 13 weeks, thus recapitulating a steatohepatitis phenotype^[32]^. In the present study, all WD-fed mice had significantly increased liver weight compared to CD-fed mice (Figure 2A). Blinded assessment of H&E-stained histological sections revealed that only WD-Cellulose feeding induced hepatic steatosis (Figure 2B&C). This finding was supported by the quantification of hepatic neutral lipids (triglycerides, free cholesterol and esterified cholesterol), which demonstrated significant lipid accumulation upon WD-feeding compared to CD-feeding regardless of the fibre source; however, the accumulation was to a lesser extent in the WD-inulin than in the WD-cellulose-fed mice (Figure 2D-F). Liver damages illustrated by plasmatic concentrations of alanine aminotransferase (ALT) and alkaline phosphatase (ALP) showed that neither the diet nor the type of dietary fibre induced hepatic damages (Figure 2G-H). Histological analysis was also performed to quantify liver inflammation and fibrosis. H&E staining revealed that neither the diet nor the type of dietary induced infiltration of inflammatory cells (Figure 2I). Finally, we assessed fibrosis by Sirius red staining, which marks type I and II collagen fibres, as well as alpha smooth muscle cell actin (α-SMA) staining, which indicates stellate cell activation in myofibroblasts (Figure 2I and 2K). We observed no significant differences in collagen deposition or stellate cell activation between the groups (Figure 2J-K).

**Figure 2:**
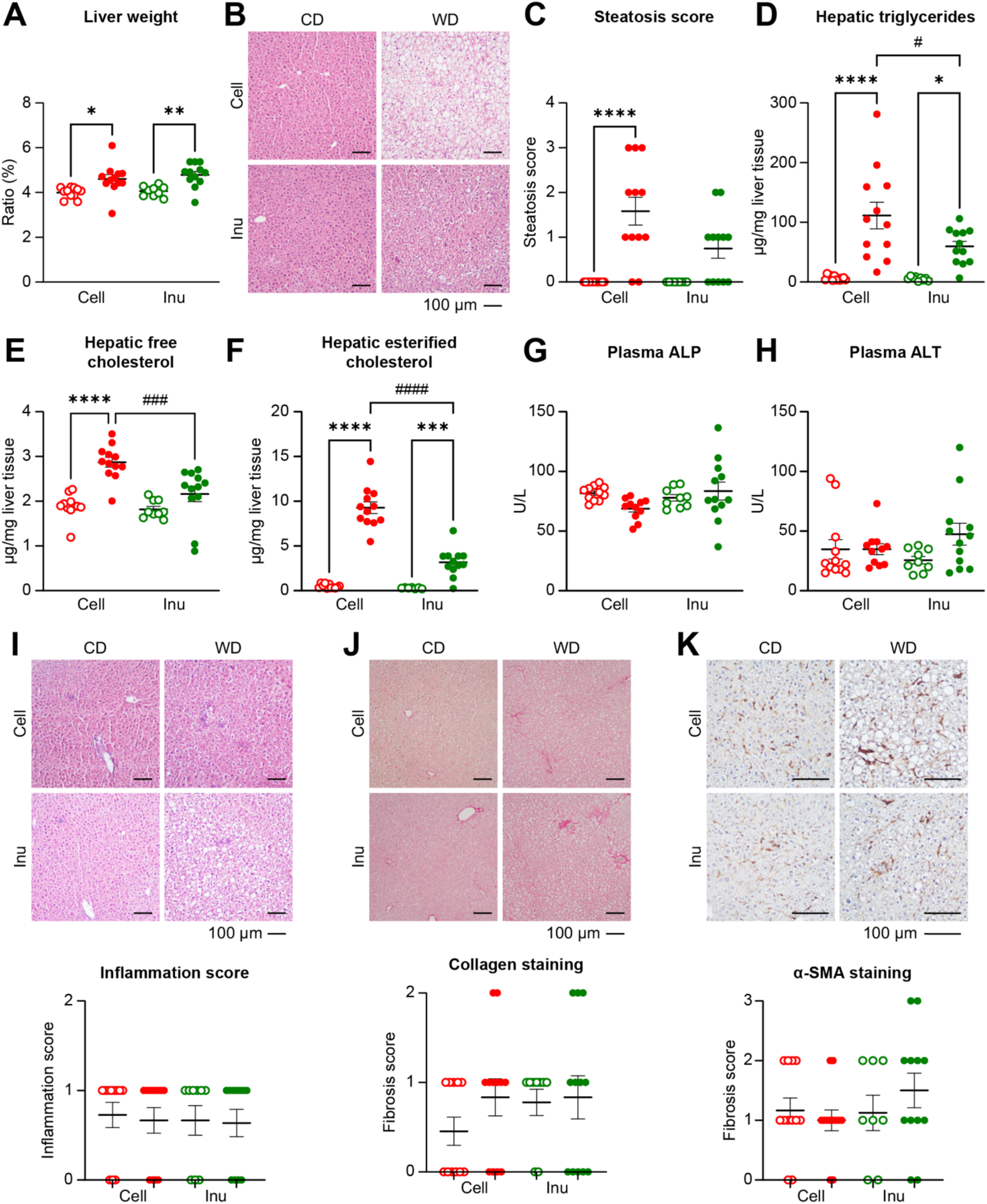
Impact of fibre-enriched WD on liver phenotype. (A) Ratio of liver to body weight (B) Representative histological sections of liver, stained with H&E, from each group at 10×. Scale bar 100 µm. (C) Steatosis score. (D-F) Hepatic neutral lipids. (G) Plasma alkaline phosphatase. (H) Plasma alanine aminotransferase. (I) Representative histological sections of liver, stained with H&E, from each group at 10×. Inflammation score. (J) Representative histological sections of liver stained with Sirius red from each group at 10x and collagen staining score. (K) Representative sections of liver staining with α-SMA from each group at 20x and α-SMA staining score. Data are presented as the mean ± SEM for n = 11-12/group.

Altogether, these results revealed that the enrichment with 9% of cellulose and inulin was protective against WD-induced steatohepatitis previously observed in mice fed with 5% of cellulose^[32]^. Moreover, the enrichment with inulin was more protective against WD-induced steatosis than that with cellulose.

### Inulin reduced WD-induced changes in hepatic gene expression

To further investigate the molecular mechanisms involved in the prevention of WD-induced MASLD associated with fibre supplementation, we analysed hepatic gene expression using microarray (Table S2). Principal component analysis (PCA) of the whole-liver transcriptome revealed a clear separation between CD-fed and WD-fed groups along the first principal component, accounting for 30.7% of the variance (Figure 3A). Interestingly, inulin-fed mice exhibited also clear separation with the cellulose-fed mice along the second principal component, accounting for 11.4% of the variance. We observed 472 differentially expressed genes (DEGs) between WD-Cellulose and CD-cellulose samples, and 331 DEGs between WD-Inulin and CD-Inulin samples (Figure 3B). The Venn diagram of these WD-induced DEGs confirmed that a majority of them were affected in WD-Cellulose mice only (Figure S2A). Hierarchical clustering of these 580 DEGs identified two clusters of genes, one containing genes with increased expression in WD-vs. CD-fed mice, independently of dietary fibre type, and one containing genes with decreased expression in WD- vs. CD-fed mice, independently of dietary fibre type (Figure S2B). These clusters were mainly implicated in lipid and xenobiotic metabolism as expected under WD (Figure S2C).

**Figure 3:**
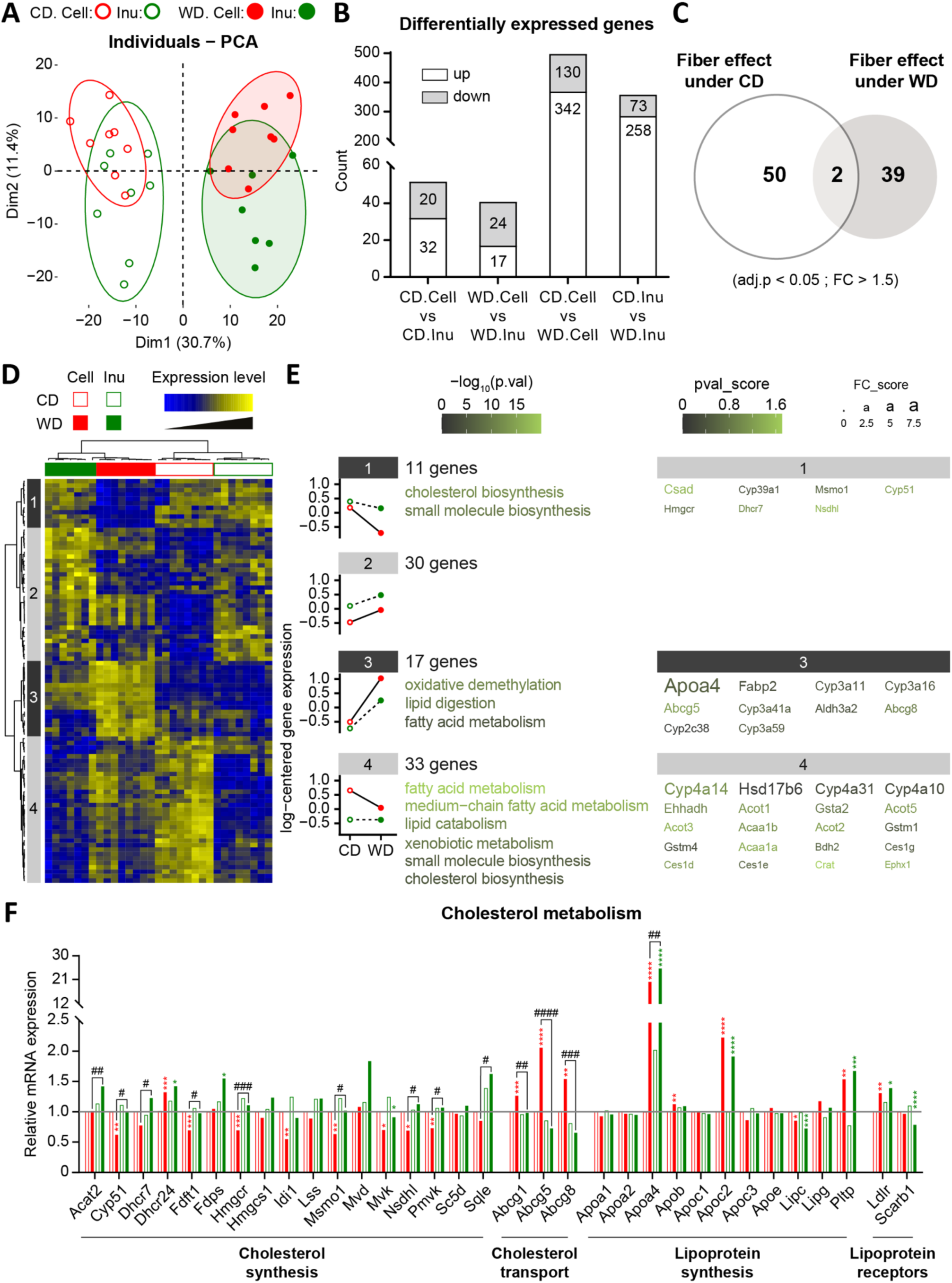
Impact of fibre-enriched WD on hepatic gene expression. (A) PCA score plots of the liver transcriptome data (n= 7/group). (B) Histogram representing the number of DEGs for each comparison. (C) Venn diagram representing the number of DEG regulated by fibre type (cellulose vs. inulin) within each diet (CD or WD). (D) Heatmap of the 91 DEGs with a significant fibre effect. (E) Mean expression profiles for the four gene clusters. The most significantly enriched biological processes and the 20 genes showing the largest differences in expression in each cluster. (F) mRNA expression of genes involved in cholesterol metabolism derived from microarray data.

We next investigated the DEGs between cellulose and inulin-fed mice in each diet. Under CD-feeding, 52 hepatic genes exhibited significantly modulated expression between cellulose and inulin, while 41 genes were differentially expressed between WD-cellulose and WD-inulin samples (Figure 3B&C). Hierarchical clustering using the 91 fibre-dependent DEGs revealed four gene clusters along the vertical axis of the heatmap (Figure 3D). In the first cluster, 11 genes exhibited lower mRNA expression among WD-fed compared to CD-fed mice, and this difference was greater under cellulose-feeding. Genes from this cluster were mostly involved in “cholesterol biosynthesis” (p = 10^-17^) and included *Msmo1*, *Cyp51*, *Hmgcr*, *Dhcr7* and *Nsdhl*. Expression of the 17 genes in cluster 3 was higher in WD-compared to CD-samples, and this difference was greater under cellulose-feeding. Genes from cluster 3 genes were associated with “lipid digestion” (p = 10^-7^) and “fatty acid metabolism” (p = 10^-2^), and included *Apoa4* and *Fabp2*, involved in lipoprotein synthesis, and *Abcg5* and *Abcg8*, involved in cholesterol transport. Finally, cluster 4 included 33 genes that exhibited decreased expression in WD-fed compared to CD-fed mice only in cellulose-supplemented mice, while no difference in expression was observed in inulin-supplemented mice. Genes from cluster 4 were involved in “fatty acid metabolism” (p = 10^-21^) and “lipid catabolism” (p = 10^-10^). Genes contributing to these pathways included *Cyp4a14*, *Cyp4a31*, *Cyp4a10* and *Ehhadh* (Figure 3E).

Then, these results led us to investigate more closely the hepatic expression of genes involved in hepatic cholesterol metabolism (Figure 3F). WD-cellulose samples had lower mRNA expression of most genes involved in cholesterol synthesis (*Cyp51*, *Fdft1*, *Hmgcr*, *Idi1*, *Msmo1*, *Mvk* and *Pmvk*) compared to CD-cellulose samples. They also displayed higher expression of cholesterol transporters (*Abcg1*, *Abcg5*, *Abcg8*) compared to CD-cellulose and WD-inulin samples. This difference in expression was not observed in WD-inulin compared to CD-inulin samples. WD-feeding also led to increased mRNA expression of apolipoproteins (*Apoa4*, *ApoB*, *ApoC2*, *Pltp*) and lipoprotein receptors (*Ldlr*, *Scarb1*) compared to CD-feeding, regardless of type of dietary fibres.

Taken together, these analyses illustrated that the enrichment of WD with inulin induced less changes in hepatic gene expression compared to enrichment with cellulose, particularly among genes involved in fatty acid metabolism and cholesterol metabolism.

### Dietary fibre impacted the fecal microbiota composition and metabolism

We next sought to describe the impact of our fibre-enriched WD on the gut microbiota. Caecum weight of WD-cellulose mice was significantly lower compared with that from CD-cellulose mice, but no change was observed between the caecum weight of WD-inulin and CD-inulin mice (Figure 4A). In addition, inulin-fed mice had significantly higher relative caecal weights compared to cellulose-fed mice, both in CD and WD-diets, confirming that inulin was fermented to a greater extent by the gut microbiota than cellulose.

**Figure 4:**
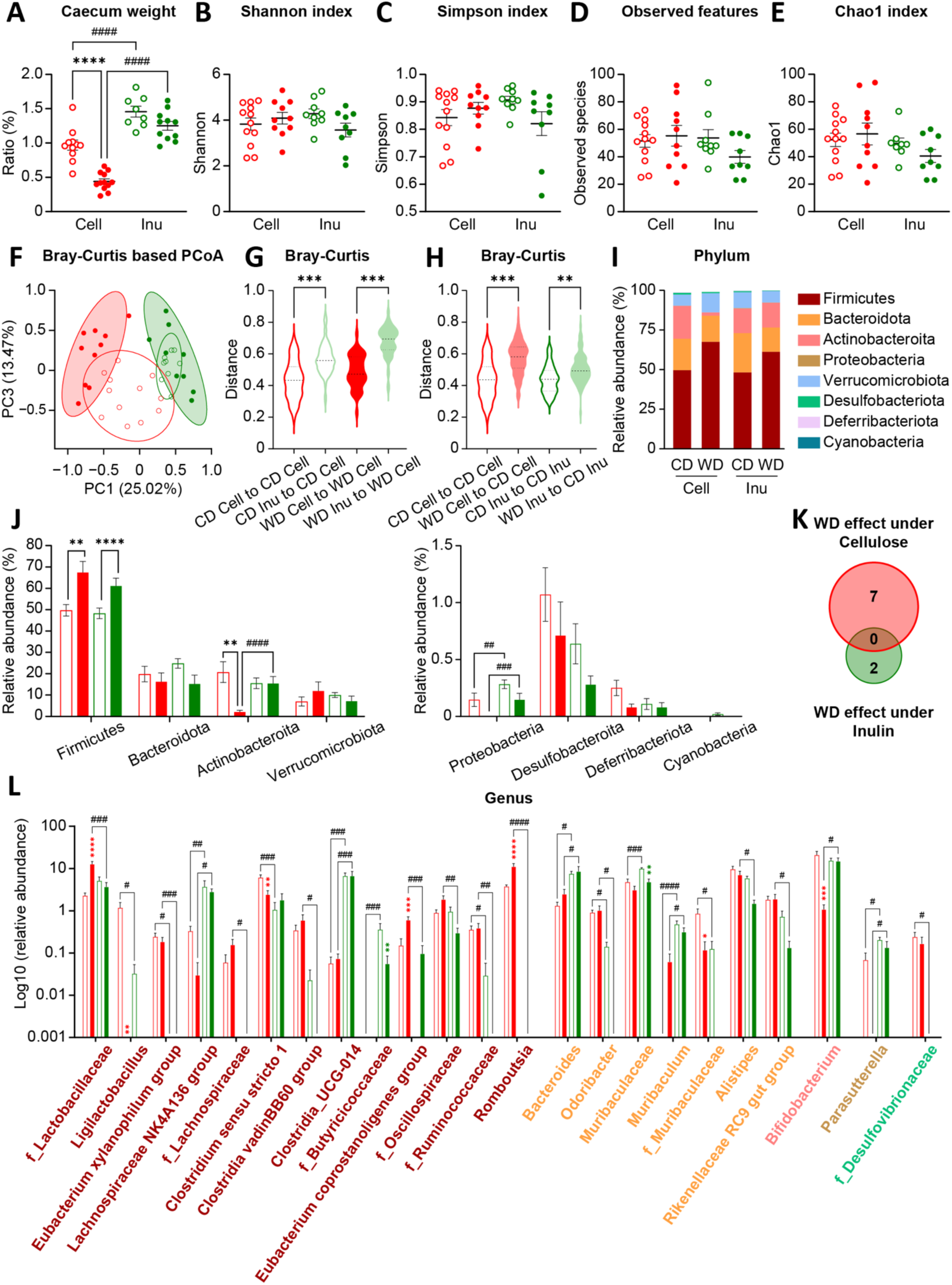
Impact of fibre-enriched WD on fecal microbiota. (A) Ratio of caecal content to body weight. (B-E) Richness (α-diversity) indices measured by Shannon index (B), Simpson index (C), observed OTU number (D) and Chao1 index (E). (F) PCoA based on the Bray-Curtis dissimilarity at the OTU level. (G-H) Bray-Curtis distance. (I-J) Relative abundance of the main phyla. (K) Venn diagram representing the number of significant genera. (L) Relative abundance of the 23 genera identified as differentially abundant between groups.

We next analysed the fecal microbiota composition using Illumina-based 16S rRNA sequencing. No difference was observed in alpha-diversity indexes between groups (Figure 4B-E). Principal coordinate analysis (PCoA) of the Bray-curtis distance showed a clear separation of the inulin groups compared to cellulose groups regardless of the diet (Figure 4F), which was confirmed by the Bray-Curtis distance (Figure 4G). Moreover, CD-inulin and WD-inulin samples clustered close together (Figure 4F), unlike CD-cellulose and WD-cellulose samples which were clearly separated from each other, this was again confirmed by the Bray-Curtis distance quantification (Figure 4H). At the phylum level, WD mice had increased abundance of *Firmicutes* regardless of the type of the dietary fibre, while only WD-cellulose mice had significant decreased abundance of *Actinobacteriota* compared to CD-cellulose mice (Figure 4I&J). Moreover, inulin fed mice had significantly higher abundance of *Proteobacteria* in both diets. At the genus level, the Venn diagram showed that 7 genera were significantly different between WD-cellulose and CD-cellulose samples, while only 2 genera were significantly different between WD-inulin and CD-inulin samples (Figure 4K). Among these 7 genera, relative abundances of *Lactobacillaceae* (family level), *Eubacterium coprostanoligenes group* and *Rombutsia* were higher, while those of *Ligilactobacillus*, *Clostridium sensu stricto 1*, *Muribaculaceae* (family level) and *Bifidobacterium* were lower, in WD-cellulose compared to CD-cellulose samples (Figure 4L). We observed that the relative abundance of *Clostridia UCG-014*, *Bacteroides* and *Muribaculum* were significantly higher in inulin groups *vs*. cellulose groups regardless of the type of diet, while the abundance of *Romboutsia* and *Lachnospiraceae* (family level) dropped drastically in inulin groups regardless of the type of diet.

Fecal metabolic profiles were obtained to assess the metabolic consequences of the observed changes in the composition of the gut microbiota. PCA of the whole ^1^H-NMR based metabolic profiles showed a clear separation between the inulin and the cellulose groups on the first principal component, illustrating that the type of dietary fibres strongly affected microbiota metabolism in both CD- and WD-fed mice. In addition, the second principal component showed a distinct clustering of CD- and WD-fed mice only under cellulose-supplemented diets, while inulin-supplemented CD- and WD- were not well discriminated (Figure S3A), which confirmed our previous observations on 16S rRNA sequencing data. Analysis of the discriminant metabolites showed no differences in fecal levels of SCFAs between the experimental groups and significant increases in many unknown aromatic metabolites in inulin-fed mice compared to cellulose-fed mice (Figure S3B). WD-cellulose fed mice displayed higher levels of all fecal bile acids detected with ^1^H-NMR compared to CD-cellulose mice, while inulin almost completely abolished these fecal bile acid concentrations for 2 of the 3 bile acids, regardless of the diet (Figure S3B). We confirmed these results with a targeted quantitative analysis of fecal bile acids using LC-MS (Figure S4). We observed significant higher levels of total and total primary bile acids in WD-cellulose vs CD-cellulose samples (Figure S4A&B). Total and total primary bile acids also tended to be higher in WD-inulin compared to CD-inulin which resulted in no significant difference between WD-cellulose and WD-inulin samples. Total secondary bile acids were higher in inulin-fed compared to cellulose-fed mice and the secondary to primary bile acid ratio was higher in CD-inuline vs CD-cellulose mice (Figure S4C&D). WD-cellulose mice had significantly higher levels of the primary bile acids αMCA, βMCA, TCA, TCDCA and TαMCA than CD-cellulose mice (Figure S4E). WD-inulin fed mice also tended to have higher levels of the most abundant primary bile acids αMCA and βMCA compared to CD-inulin fed mice. These modifications in primary and secondary fecal bile acids led us to analyse the hepatic expression of genes involved in bile acid metabolism. Neither the diet nor the type of dietary fibre altered the hepatic expression of genes involved in bile acid synthesis (*Cyp7a1*, *Cyp27a1*, *Cyp7b1*, *Cyp8b1*) (Figure S5).

Collectively, these results show that both the type of dietary fibres and other dietary components from the WD affect the intestinal microbiota composition and metabolism, with a potential interaction of both factors.

### Correlations between metabolic parameters and microbiota alterations

To gain insights between the potential relationships between fibre-induced metabolic outcomes and gut microbiota genera, we performed correlation analyses (Figure 5A&B). We observed many significant correlations between several genera and plasma triglyceride levels. Two bacterial genera were correlated with almost all metabolic parameters: *Bifidobacterium* were inversely related to plasma lipids (triglycerides, total cholesterol, HDL and LDL), to weight-related parameters (body weight gain and subcutaneous WAT weight) and to liver steatosis markers (triglycerides and cholesterol esters); while *Eubacterium_coprostanoligenes* were positively related to all plasma lipid levels, body weight parameters and liver steatosis markers. Using another multivariate correlation approach that better handles high-dimensional data, we confirmed the strong positive relationship between fecal abundance of *Eubacterium_coprostanoligenes* and diet-induced metabolic perturbations, and also highlighted similar correlations with *Rombutsia* and *Lactobaccilaceae* (family level).

**Figure 5:**
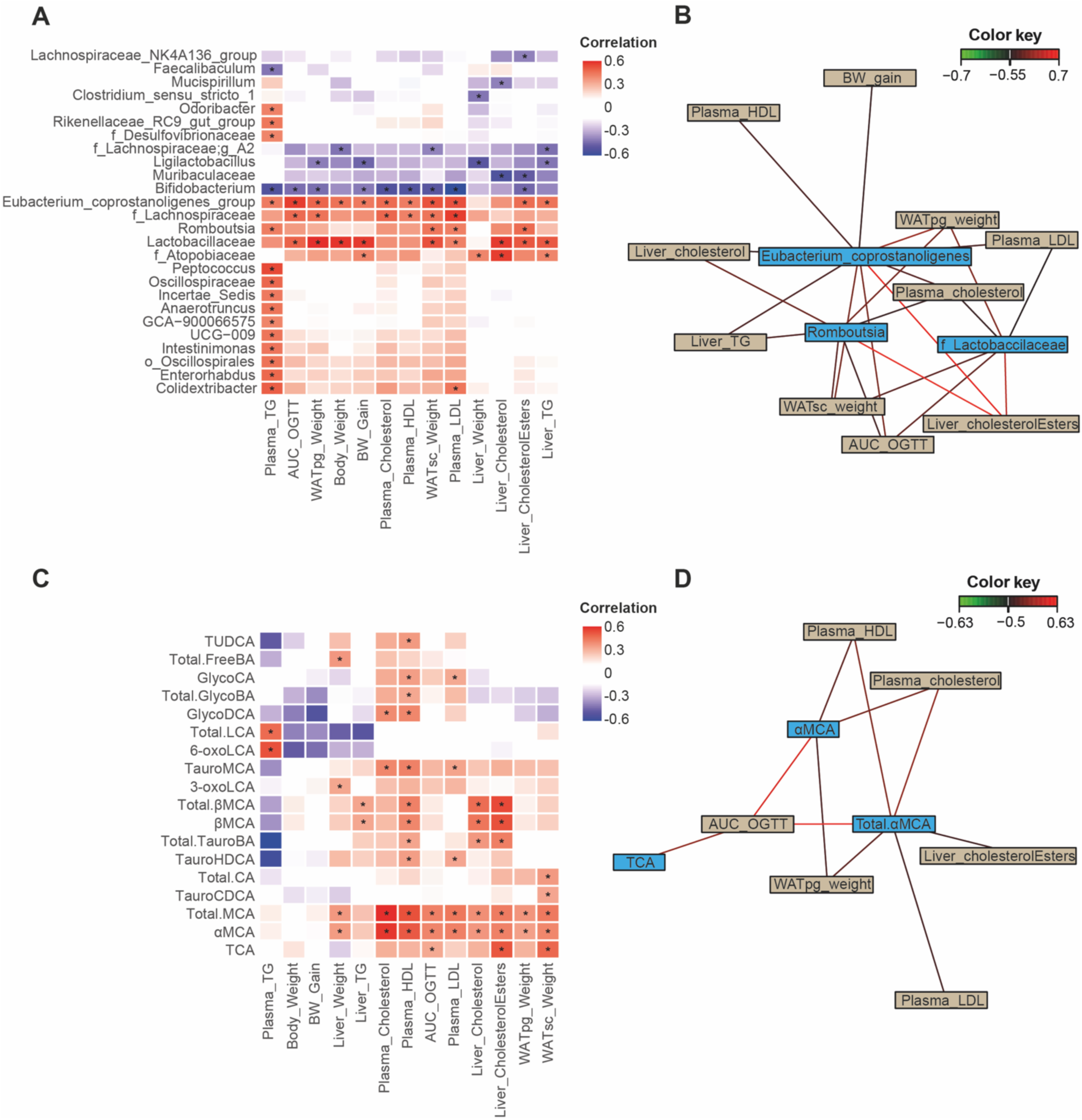
Correlations between metabolic parameters and fecal markers of gut microbiota changes. (A) Heatmap of Spearman’s correlations between microbial genera and metabolic parameters. (B) Relevance network of bacterial genera and metabolic parameters selected with rCCA. (C) Heatmap of Spearman’s correlations between fecal bile acids and metabolic parameters. (D) Relevance network of fecal bile acids and metabolic parameters selected with rCCA. *: p < 0.01 for spearman’s correlation. For relevance networks, only associations with and absolute correlation > 0.5 are represented.

We performed the same correlations between metabolic parameters and fecal bile acids and found the strongest positive correlation between MCA levels (both total MCA and αMCA) and plasma cholesterol levels (R^2^=0.6). More generally, total MCA and αMCA levels displayed the highest number of significant correlations with the metabolic outcomes affected by the diets. Multivariate correlation analysis confirmed this result and more precisely highlighted the positive relationships between total MCA and αMCA and markers of cholesterol homeostasis, namely plasma HDL, LDL and total cholesterol, and liver cholesterol esters.

## Discussion

MASLD has become the most prevalent chronic liver disease worldwide, with a 30% mean prevalence in adult population^[1,39]^. Since diet-induced obesity is a major risk factor driving MASLD development, dietary modification is a promising strategy for ameliorating MASLD and its progression to MASH. Several studies have shown that consumption of plant-based diet naturally abundant in fermentable soluble fibre, such as inulin, is associated with numerous health benefits, including improvements of metabolic parameters and reduction in obesity^[24,40,41]^. However, little is known about the differential effects of soluble vs. insoluble dietary fibres on liver metabolism. Previous studies combining diet-induced metabolic perturbations with dietary fibres have mostly used lipid-enriched diets (HFD)^[40–42]^, sucrose-enriched diets^[43]^, high fructose diets^[44]^, or choline-deficient high-fat diet^[45]^, which might not represent the most appropriate dietary models to study MASLD development. Here, we compared the metabolic and hepatic effects of a soluble fibre (inulin) and an insoluble fibre (cellulose) in thermoneutral housed, WD-fed mice.

We found that inulin enrichment alleviated WD-induced obesity (body weight gain and subcutaneous and perigonadic WAT), glucose intolerance and dyslipidaemia compared to cellulose enrichment. These results are consistent with many previous studies showing that inulin significantly decreased metabolic perturbations induced by high fat, high sucrose or high fructose diets^[40,42–44]^. Nevertheless, two others studies have reported no significant differences in obesity and glucose tolerance between HFD-fed mice with inulin or cellulose^[41,46]^. This discrepancy might be due to differences in the duration of the dietary interventions (6 weeks in^[41]^ or 10 weeks in^[46]^ vs 18 weeks in our study), or in the type and proportion of fibre used. The mechanisms allowing inulin to modulate weight are not fully understood but are thought to involve regulation of satiety through its capacity to form a viscous gel and to modulate secretion of gastrointestinal hormones signalling to the brain^[23,47]^. However, in our study, WD-inulin-fed mice consumed more calories than WD-cellulose-fed, thus differential effect on satiety is probably not involved. Other possible mechanisms include reduced intestinal lipid absorption, improved gut barrier function^[48,49]^ and the induction of intestinal gluconeogenesis by SCFAs produced by inulin^[50]^. Intestinal gluconeogenesis has been shown to promote energy expenditure and reduce hepatic steatosis^[51]^.

Regarding MASLD, we first observed that enriching WD with 9% dietary fibres (both cellulose and inulin) induced significant hepatic steatosis (triglyceride accumulation), but the mice did not develop a significant MASH phenotype (no inflammation and no fibrosis), in both cellulose- and inulin-enrichment conditions. This represents in itself an interesting finding since our previous studies^[29,32]^ and those of others^[30]^ using WD enriched with the same amount of lipids (≈20%), sucrose (≈30%) and cholesterol (0.2%), for similar durations (15 to 18 weeks), in male mice from the same genetic background, have consistently observed MASH development, which was further aggravated by housing in thermoneutral conditions^[32]^, as performed in the present study. Thus, we could postulate that WD-enrichment with 9% fibre was sufficient to protect mice from MASH, and that this protection was seen with both cellulose and inulin. Indeed, commercial WD used in previous studies usually contains only 4-5% of fibre^[29,30,32]^. This result is in line with a previous study in mice fed a HFD or a HFD+10% inulin demonstrating that inulin supplementation decreased hepatic steatosis, inflammatory gene expression and immune cell populations^[52]^, and with many studies showing a dose-dependency in fibre-induced metabolic benefits both in humans^[53]^ and in rodents^[54]^. However, there could also be other compositional differences between the WD used in the present study and those used in previous studies explaining the difference in the extend of MASH development.

Next, we found that inulin was more protective against WD-induced steatosis than cellulose. Consistently, inulin-enriched WD induced less changes in hepatic gene expression than cellulose-enriched WD, especially for genes involved in lipid and cholesterol metabolism. Soluble inulin was previously shown to be more efficient than insoluble cellulose to reduce hepatic steatosis, necro-inflammation, ballooning and fibrosis in a choline-deficient high-fat diet model of MASLD^[45]^. In HFD models of MASLD, effects of inulin on hepatic steatosis are more controversial and only a slight decrease of hepatic triglyceride contents was observed^[41,55]^, with no effect observed on hepatic cholesterol. Using a WD in the present study allowed us to unravel a significant effect of inulin on hepatic cholesterol metabolism. At the gene level, our results demonstrated that WD-cellulose decreased mRNA expression of most genes involved in cholesterol synthesis compared to CD-cellulose. These results are consistent with literature since when dietary cholesterol intake increases, a feedback mechanism decreases hepatic expression of genes involved in cholesterol synthesis^[56]^. However, this difference in cholesterol gene expression was not seen between WD-inulin and CD-inulin samples. Moreover, we observed that levels of plasma HDL-, LDL, total cholesterol and hepatic free and esterified cholesterol were significantly less increased upon WD in inulin mice than in cellulose mice. It is widely accepted that soluble fibre act primarily in the intestine to reduce dietary fat and cholesterol uptake, thus promoting secondary responses in the liver and peripheral circulation^[57,58]^. These effects of dietary inulin on reduction of intestinal cholesterol absorption have been reported in both human subjects^[59]^, animal models^[60]^ and *in vitro* study on rat proximal small intestine^[61]^. Although the exact molecular mechanisms have not been clearly defined, they are most often attributed to the viscosity of soluble fibres, which interfere with cholesterol absorptive process. Soluble fibres can indeed directly bind cholesterol within the intestinal lumen^[62]^, delay the diffusion of cholesterol from the lumen to gut epithelial cells^[57]^, and/or reduce biliary emulsification of cholesterol^[63]^.

The beneficial metabolic effects of inulin have been repeatedly shown to depend on its impact on the gut microbiota and its fermentation to beneficial metabolites including SCFAs^[24,52,64]^. Here, we observed significant differences in microbiota composition (with 16S rRNA sequencing) and metabolism (with untargeted NMR-based metabolomics) between inulin- and cellulose-fed mice, independently of the diet composition. As reflected by unsupervised analysis, fecal metabolic profiles were even more affected by the type of fibre than by the type of diet. Moreover, we observed that WD-induced perturbations in microbiota were less pronounced in inulin-fed than in cellulose-fed mice, which could reflect a higher resilience of the inulin-selected gut microbiota toward WD. At the phylum level, cellulose-enriched WD decreased *Actinobacteroita* abundance, which was mitigated in inulin-WD. At the genus level, WD-cellulose decreased beneficial *Bifidobacterium* and increased *Lactobacillaceae* (family level), while inulin prevented these changes. It has previously been described that inulin prevents gut microbiota dysbiosis induced by HFD^[42]^. Independently of the diets, we also observed that inulin completely abolished the abundance of *Desulfovibrionaceae*, which was previously observed in a mouse study with different sources of inulin^[41]^ and is consistent with studies showing that sulfate-reducing bacteria were positively correlated with obesity^[65]^. Similarly, inulin is known to favour the growth of SCFA-producing bacteria, such as *Bifidobacterium*, *Butyricicoccaceae*, *Parasutterella*, which we found higher in inulin-fed mice than in cellulose-fed mice. These bacteria have been negatively correlated with metabolic perturbations (BMI, serum lipids, blood glucose…) in obese adults^[66]^. However, measuring SCFAs in feces, we did not detect significant variation between groups, which does not necessarily reflect no difference in SCFA production by bacteria, since they can be absorbed or metabolized. Although SCFAs are the primary metabolic products of soluble fibre fermentation, other metabolites can be produced directly or indirectly upon inulin metabolism or their concentrations can be indirectly influenced during this process^[23]^. Among these, bile acids are not directly produced by dietary fibre fermentation but their profiles are affected. In our study, we indeed observed using untargeted metabolomics that bile acids were among the most different metabolites between inulin- and cellulose-diets. Another study using untargeted metabolomics in blood and fecal samples of inulin vs cellulose fed mice also found that the largest and most abundant family of differentially abundant metabolites was represented by bile acids^[67]^. Here, we thus conducted a comprehensive targeted analysis of fecal bile acids and observed significant increase in the total and total primary bile acid levels in WD-cellulose vs CD-cellulose samples. WD also tended to increase total and total primary bile acids in inulin fed mice and there was no significant difference between WD-cellulose and WD-inulin samples. However, this WD-dependant increase was probably not due to increased hepatic bile acid synthesis as there was no change in the hepatic expression of genes involved in bile acid synthesis, metabolism and transport. Total secondary bile acids were increased in inulin-fed compared to cellulose-fed mice and the secondary to primary bile acid ratio was higher in CD-inuline vs CD-cellulose mice, which probably reflects the inulin-driven enrichement of specific bacterial taxa possessing bile salt hydrolase or 7α-dehydroxylase activity. At the individual bile acid level, most statistically significant changes were observed between WD-cellulose and CD-cellulose groups with increased levels of the highly abundant αMCA and βMCA and their tauro-conjugated forms. WD also tended to increase these bile acids in inulin fed mice and there was no significant difference between WD-cellulose and WD-inulin samples. Most studies that investigated the impact of inulin on bile acid metabolism have compared inulin-enriched diets to diets containing a much lower percentage of fibre^[67–70]^ and the results regarding fecal bile acids are contradictory. Some studies report decreased levels of fecal bile acids upon inulin supplementation^[67,68]^ and postulate enhanced bile acid reabsorption, but these studies are conducted in a context of diets not enriched in lipids. In contrast, others studies have found higher levels of total bile acids in animals fed HFD^[69]^ or high fat high sucrose diets^[70]^ enriched with inulin.

Finally, we calculated correlations between changes observed in gut microbiota composition and in metabolic parameters and observed a positive correlation between the abundance of *Eubacterium coprostanoligenes* and most of the metabolic parameters including hepatic levels of cholesterol esters and plasma cholesterol. More precisely, this bacteria was increased upon WD in cellulose fed mice and strongly decreased upon inulin. This finding was quite surprising as *E. coprostanoligenes* is usually considered as a beneficial bacteria as it was negatively correlated to body mass index^[71,72]^, increased with inulin supplementation in obese children^[72]^ and decreased in various models of diet-induced obesity in rodents^[73,74]^. *E. coprostanoligenes* is well known for its ability to reduce cholesterol to coprostanol. Coprostanol exhibits lower intestinal absorption than cholesterol and therefore increased fecal elimination in human^[75]^. Increasing cholesterol-to-coprostanol conversion has thus been suggested as a strategy to lower circulating cholesterol levels in the host^[76]^. Several studies with animal models were designed to investigate the effect of feeding of *E. coprostanoligenes* on serum cholesterol concentration but showed contradictory results with cholesterol lowering effects in rabbits^[77]^ and no effect observed in laying hens^[78]^ or germ-free mice^[78]^. The mechanisms linking *E. coprostanoligenes* to systemic and hepatic cholesterol metabolism in our experiment remain to be clarified.

In summary, our findings are in agreement with a model that explains superior beneficial metabolic effects of soluble fibres type inulin over those induced by insoluble fibres type cellulose on MASLD development, which may involve the gut microbiota-liver axis. These beneficial effects include protection against WD-induced body weight gain, hyperglycemic response, hepatic steatosis and hepatic cholesterol metabolism perturbations. Protection was observed for both types of fibre but was of higher magnitude with inulin. Of note, both cellulose and inulin were associated with the absence of WD-induced MASH. WD-induced perturbations of gut microbiota composition and metabolism were also lower upon inulin enrichment, suggesting that inulin-selected bacterial taxa may be less affected by dietary components of the WD. Our data confirm the relevance of increasing dietary fibre intake to alleviate MASLD development.

## Limitations of the study

A main limitation of our study is the exclusive use of male mice, despite the sexually dimorphic nature of MASLD^[29]^ and the known sex-specific differences in gut microbiota composition^[79]^. Additionally, our 18-week intervention, although sufficient to observe significant metabolic and hepatic changes, may not fully capture the long-term effects of dietary fibre supplementation on MASLD progression to MASH. The use of a single fibre concentration (9% total dietary fibre) limits our understanding of dose-response relationships, and the specific inulin formulation used may not represent the diverse range of soluble fibres available in human diets. Furthermore, while we observed significant changes in fecal bile acid metabolism and gut microbiota composition, the mechanistic pathways linking these changes to hepatic protection require further investigation through targeted functional studies.

## Author contributions

Conceptualization: N.L. and S.E.S.; Formal analysis: C.M.P.M., A.P., V.A.B., B.C., Y.L., J.B.M., A.G.S., X.B., H.G., A.F., N.L. and S.E.S.; Investigation: C.M.P.M., M.M., A.P., V.A.B., J.G., F.L., P.P., M.H., J.B., A.F., L.G.P., B.C., C.D., E.R.B., Y.L., C.N., N.G., A.D., J.B.M., C.C., A.D., A.G.S., X.B., A.T., H.G., N.L. and S.E.S.; Writing—original draft preparation: C.M.P.M., S.E.S. and N.L.; Writing—review and editing, S.E.S, N.L., C.M.P.M. and H.G.; Visualization: C.M.P.M., A.P., S.E.S., and N.L.; Supervision: S.E.S, N.L., and H.G.; Funding acquisition: N.L., S.E.S. and H.G. All authors have read and agreed to the published version of the manuscript.

## Acknowledgements

This work was supported by the French Foundation for the Medical Research FRM (Equipe FRM EQU202303016327). This work was also supported by grant from the French National Research Agency (ANR) GADGET (ANR-22-CE34-0005). We thank Anexplo (Genotoul, Toulouse), We-Met facility (Inserm U1297, Toulouse) for their excellent work on plasma biochemistry.

## References

[1] Z. M. Younossi, P. Golabi, J. M. Paik, A. Henry, C. Van Dongen, L. Henry, Hepatology, 2023, 77, 1335–1347.

[2] M. Eslam, A. J. Sanyal, J. George, A. Sanyal, B. Neuschwander-Tetri, C. Tiribelli, D. E. Kleiner, E. Brunt, E. Bugianesi, H. Yki-Järvinen, H. Grønbæk, H. Cortez-Pinto, J. George, J. Fan, L. Valenti, M. Abdelmalek, M. Romero-Gomez, M. Rinella, M. Arrese, M. Eslam, P. Bedossa, P. N. Newsome, Q. M. Anstee, R. Jalan, R. Bataller, R. Loomba, S. Sookoian, S. K. Sarin, S. Harrison, T. Kawaguchi, V. W.-S. Wong, V. Ratziu, Y. Yilmaz, Z. Younossi, Gastroenterology, 2020, 158, 1999–2014.e1.

[3] M. Eslam, P. N. Newsome, S. K. Sarin, Q. M. Anstee, G. Targher, M. Romero-Gomez, S. Zelber-Sagi, V. Wai-Sun Wong, J.-F. Dufour, J. M. Schattenberg, T. Kawaguchi, M. Arrese, L. Valenti, G. Shiha, C. Tiribelli, H. Yki-Järvinen, J.-G. Fan, H. Grønbæk, Y. Yilmaz, H. Cortez-Pinto, C. P. Oliveira, P. Bedossa, L. A. Adams, M.-H. Zheng, Y. Fouad, W.-K. Chan, N. Mendez-Sanchez, S. H. Ahn, L. Castera, E. Bugianesi, V. Ratziu, J. George, J. Hepatol., 2020, 73, 202–209.

[4] M. E. Rinella, B. A. Neuschwander-Tetri, M. S. Siddiqui, M. F. Abdelmalek, S. Caldwell, D. Barb, D. E. Kleiner, R. Loomba, Hepatology, 2023, 77, 1797–1835.

[5] J. C. Cohen, J. D. Horton, H. H. Hobbs, Science (80-.). 2011, 332, 1519–1523.

[6] A. Fougerat, A. Montagner, N. Loiseau, H. Guillou, W. Wahli, Cells, 2020, 9, 1638.

[7] E. E. Powell, V. W.-S. Wong, M. Rinella, Lancet, 2021, 397, 2212–2224.

[8] European Association for the Study of the Liver, European Association for the Study of Diabetes, European Association for the Study of Obesity, Diabetologia 2024, 67, 2375–2392.

[9] Y. Xiao, X. Zhang, D. Yi, F. Qiu, L. Wu, Y. Tang, N. Wang, Front. Nutr., 2023, 10, 1225946.

[10] S. V Thompson, B. A. Hannon, R. An, H. D. Holscher, Am. J. Clin. Nutr., 2017, 106, 1514–1528.

[11] A. N. Reynolds, A. Akerman, S. Kumar, H. T. Diep Pham, S. Coffey, J. Mann, BMC Med., 2022, 20, 139.

[12] H. Colak, G. N. F. Larik, M. A. van Baak, E. E. Canfora, Clin. Nutr., 2025, 52, 236–251.

[13] V. A. Arita, M. C. Cabezas, J. A. Hernández Vargas, S. J. Trujillo-Cáceres, N. Mendez Pernicone, L. A. Bridge, H. Raeisi-Dehkordi, C. A. W. Dietvorst, R. Dekker, J. P. Uriza-Pinzón, M. Tawfik, K. A. Berk, J. Massoels, S. Driessen, M. E. Tushuizen, A. G. Holleboom, D. E. Grobbee, O. H. Franco, S. Beigrezaei, GRIPonMASH Consortium, BMC Med., 2025, 23, 502.

[14] A. A. Sangouni, S. Hassani Zadeh, H. Mozaffari-Khosravi, M. Hosseinzadeh, Br. J. Nutr., 2022, 128, 1231–1239.

[15] L. Haigh, C. Kirk, K. El Gendy, J. Gallacher, L. Errington, J. C. Mathers, Q. M. Anstee, Clin. Nutr., 2022, 41, 1913–1931.

[16] Y. Xiong, X. Shi, X. Xiong, S. Li, H. Zhao, H. Song, J. Wang, L. Zhang, S. You, G. Ji, B. Liu, N. Wu, Food Funct., 2024, 15, 8248–8257.

[17] Y. Ni, L. Qian, S. L. Siliceo, X. Long, E. Nychas, Y. Liu, M. J. Ismaiah, H. Leung, L. Zhang, Q. Gao, Q. Wu, Y. Zhang, X. Jia, S. Liu, R. Yuan, L. Zhou, X. Wang, Q. Li, Y. Zhao, H. El-Nezami, A. Xu, G. Xu, H. Li, G. Panagiotou, W. Jia, Cell Metab., 2023, 35, 1530–1547.e8.

[18] A. Tripathi, J. Debelius, D. A. Brenner, M. Karin, R. Loomba, B. Schnabl, R. Knight, Nat. Rev. Gastroenterol. Hepatol., 2018, 15, 397–411.

[19] J. Aron-Wisnewsky, M. V. Warmbrunn, M. Nieuwdorp, K. Clément, Gastroenterology 2020, 158, 1881–1898.

[20] M. R. Bomhof, J. A. Parnell, H. R. Ramay, P. Crotty, K. P. Rioux, C. S. Probert, S. Jayakumar, M. Raman, R. A. Reimer, Eur. J. Nutr., 2019, 58, 1735–1745.

[21] F. Bakhshimoghaddam, K. Shateri, M. Sina, M. Hashemian, M. Alizadeh, J. Nutr., 2018, 148, 1276–1284.

[22] V. V. Huwiler, K. A. Schönenberger, A. Segesser von Brunegg, E. Reber, S. Mühlebach, Z. Stanga, M. L. Balmer, Nutrients, 2022, 14, 2627.

[23] N. M. Delzenne, L. B. Bindels, A. M. Neyrinck, J. Walter, Nat. Rev. Microbiol., 2025, 23, 225–238.

[24] B. Chassaing, J. Miles-Brown, M. Pellizzon, E. Ulman, M. Ricci, L. Zhang, A. D. Patterson, M. Vijay-Kumar, A. T. Gewirtz, Am. J. Physiol. Liver Physiol., 2015, 309, G528–G541.

[25] T. M. Barber, S. Kabisch, A. F. H. Pfeiffer, M. O. Weickert, Nutrients, 2020, 12, 3209.

[26] J. S. L. de Munter, F. B. Hu, D. Spiegelman, M. Franz, R. M. van Dam, PLoS Med., 2007, 4, e261.

[27] M. B. Schulze, M. Schulz, C. Heidemann, A. Schienkiewitz, K. Hoffmann, H. Boeing, Arch. Intern. Med., 2007, 167, 956–65.

[28] M. O. Weickert, A. F. H. Pfeiffer, J. Nutr., 2008, 138, 439–42.

[29] S. Smati, A. Polizzi, A. Fougerat, S. Ellero-Simatos, Y. Blum, Y. Lippi, M. Régnier, A. Laroyenne, M. Huillet, M. Arif, C. Zhang, F. Lasserre, A. Marrot, T. Al Saati, J. Wan, C. Sommer, C. Naylies, A. Batut, C. Lukowicz, T. Fougeray, B. Tramunt, P. Dubot, L. Smith, J. Bertrand-Michel, N. Hennuyer, J. P. Pradere, B. Staels, R. Burcelin, F. Lenfant, J. F. Arnal, T. Levade, L. Gamet-Payrastre, S. Lagarrigue, N. Loiseau, S. Lotersztajn, C. Postic, W. Wahli, C. Bureau, M. Guillaume, A. Mardinoglu, A. Montagner, P. Gourdy, H. Guillou, Gut, 2022, 71, 807–821.

[30] M. Vacca, I. Kamzolas, L. M. Harder, F. Oakley, C. Trautwein, M. Hatting, T. Ross, B. Bernardo, A. Oldenburger, S. T. Hjuler, I. Ksiazek, D. Lindén, D. Schuppan, S. Rodriguez-Cuenca, M. M. Tonini, T. R. Castañeda, A. Kannt, C. M. P. Rodrigues, S. Cockell, O. Govaere, A. K. Daly, M. Allison, K. Honnens de Lichtenberg, Y. O. Kim, A. Lindblom, S. Oldham, A.-C. Andréasson, F. Schlerman, J. Marioneaux, A. Sanyal, M. B. Afonso, R. Younes, Y. Amano, S. L. Friedman, S. Wang, D. Bhattacharya, E. Simon, V. Paradis, A. Burt, I. M. Grypari, S. Davies, A. Driessen, H. Yashiro, S. Pors, M. Worm Andersen, M. Feigh, C. Yunis, P. Bedossa, M. Stewart, H. L. Cater, S. Wells, J. M. Schattenberg, Q. M. Anstee, LITMUS Investigators, D. Tiniakos, J. W. Perfield, E. Petsalaki, P. Davidsen, A. Vidal-Puig, Nat. Metab., 2024, 6, 1178–1196.

[31] S. Gallage, J. E. B. Avila, P. Ramadori, E. Focaccia, M. Rahbari, A. Ali, N. P. Malek, Q. M. Anstee, M. Heikenwalder, Nat. Metab., 2022, 4, 1632–1649.

[32] C. M. P. Martin, A. Polizzi, V. Alquier-Bacquié, M. Huillet, C. Rives, C. Dauriat, J. Bruse, V. Melin, C. Naylies, Y. Lippi, F. Lasserre, J. Wan, R. Flores-Flores, J. Bertrand-Michel, F. Blas-Y-Estrada, E. Rousseau-Bacquié, T. Levade, H. Rémignon, D. Langin, E. Mouisel, S. Lotersztajn, B. Chassaing, L. Gamet-Payrastre, H. Guillou, S. Ellero-Simatos, A. Fougerat, N. Loiseau, iScience, 2025, 113221.

[33] D. E. Kleiner, E. M. Brunt, M. Van Natta, C. Behling, M. J. Contos, O. W. Cummings, L. D. Ferrell, Y.-C. Liu, M. S. Torbenson, A. Unalp-Arida, M. Yeh, A. J. McCullough, A. J. Sanyal, Nonalcoholic Steatohepatitis Clinical Research Network, Hepatology, 2005, 41, 1313–21.

[34] C. Rives, C. M. P. Martin, L. Evariste, A. Polizzi, M. Huillet, F. Lasserre, V. Alquier-Bacquie, P. Perrier, J. Gomez, Y. Lippi, C. Naylies, T. Levade, F. Sabourdy, H. Remignon, P. Fafournoux, B. Chassaing, N. Loiseau, H. Guillou, S. Ellero-Simatos, L. Gamet-Payrastre, A. Fougerat, Mol. Nutr. Food Res., 2024, 68, e2300491.

[35] E. Bolyen, J. R. Rideout, M. R. Dillon, N. A. Bokulich, C. C. Abnet, G. A. Al-Ghalith, H. Alexander, E. J. Alm, M. Arumugam, F. Asnicar, Y. Bai, J. E. Bisanz, K. Bittinger, A. Brejnrod, C. J. Brislawn, C. T. Brown, B. J. Callahan, A. M. Caraballo-Rodríguez, J. Chase, E. K. Cope, R. Da Silva, C. Diener, P. C. Dorrestein, G. M. Douglas, D. M. Durall, C. Duvallet, C. F. Edwardson, M. Ernst, M. Estaki, J. Fouquier, J. M. Gauglitz, S. M. Gibbons, D. L. Gibson, A. Gonzalez, K. Gorlick, J. Guo, B. Hillmann, S. Holmes, H. Holste, C. Huttenhower, G. A. Huttley, S. Janssen, A. K. Jarmusch, L. Jiang, B. D. Kaehler, K. Bin Kang, C. R. Keefe, P. Keim, S. T. Kelley, D. Knights, I. Koester, T. Kosciolek, J. Kreps, M. G. I. Langille, J. Lee, R. Ley, Y.-X. Liu, E. Loftfield, C. Lozupone, M. Maher, C. Marotz, B. D. Martin, D. McDonald, L. J. McIver, A. V Melnik, J. L. Metcalf, S. C. Morgan, J. T. Morton, A. T. Naimey, J. A. Navas-Molina, L. F. Nothias, S. B. Orchanian, T. Pearson, S. L. Peoples, D. Petras, M. L. Preuss, E. Pruesse, L. B. Rasmussen, A. Rivers, M. S. Robeson, P. Rosenthal, N. Segata, M. Shaffer, A. Shiffer, R. Sinha, S. J. Song, J. R. Spear, A. D. Swafford, L. R. Thompson, P. J. Torres, P. Trinh, A. Tripathi, P. J. Turnbaugh, S. Ul-Hasan, J. J. J. van der Hooft, F. Vargas, Y. Vázquez-Baeza, E. Vogtmann, M. von Hippel, W. Walters, Y. Wan, M. Wang, J. Warren, K. C. Weber, C. H. D. Williamson, A. D. Willis, Z. Z. Xu, J. R. Zaneveld, Y. Zhang, Q. Zhu, R. Knight, J. G. Caporaso, Nat. Biotechnol., 2019, 37, 852–857.

[36] H. Lin, S. Das Peddada, Nat. Commun., 2020, 11, 3514.

[37] C. Lukowicz, S. E. Simatos, M. Régnier, A. Polizzi, F. Lasserre, A. Montagner, Y. Lippi, E. L. Jamin, J. F. Martin, C. Naylies, C. Canlet, L. Debrauwer, J. Bertrand-Michel, T. Al Saati, V. Théodorou, N. Loiseau, L. M. Lakhal, H. Guillou, L. Gamet-Payrastre, Environ. Health Perspect., 2018, 126, 1–18.

[38] F. Rohart, B. Gautier, A. Singh, K.-A. Lê Cao, PLoS Comput. Biol., 2017, 13, e1005752.

[39] L. Miao, G. Targher, C. D. Byrne, Y. Y. Cao, M. H. Zheng, Trends Endocrinol. Metab., 2024, 35, 697–707.

[40] J. Zou, B. Chassaing, V. Singh, M. Pellizzon, M. Ricci, M. D. Fythe, M. V. Kumar, A. T. Gewirtz, Cell Host Microbe, 2018, 23, 41–53.e4.

[41] A. M. Neyrinck, J. Rodriguez, C. R. Sánchez, M. Autuori, P. D. Cani, L. B. Bindels, J. Bindelle, N. M. Delzenne, Eur. J. Nutr., 2025, 64.

[42] M. Albouery, A. Bretin, B. Buteau, S. Grégoire, L. Martine, S. Gambert, A. M. Bron, N. Acar, B. Chassaing, M. A. Bringer, Nutrients, 2021, 13.

[43] Z. Yang, H. Su, Y. Lv, H. Tao, Y. Jiang, Z. Ni, L. Peng, X. Chen, Food Res. Int., 2023, 163, 112309.

[44] S. Jung, H. Bae, W. S. Song, Y. Chun, J. Le, Y. Alam, A. Verlande, S. K. Chun, J. Kim, M. E. Kelly, M. L. Lopez, S. H. Park, D. Onofre, J. Baek, K. H. Jang, V. I. Rubtsova, A. Anica, S. Masri, G. Lee, C. Jang, Nat. Metab., 2025.

[45] W. Wei, C. C. Wong, Z. Jia, W. Liu, C. Liu, F. Ji, Y. Pan, F. Wang, G. Wang, L. Zhao, E. S. H. Chu, X. Zhang, J. J. Y. Sung, J. Yu, Nat. Microbiol., 2023, 8, 1534–1548.

[46] L. L. Li, Y. T. Wang, L. M. Zhu, Z. Y. Liu, C. Q. Ye, S. Qin, Sci. Rep., 2020, 10, 1–12.

[47] M. Akhlaghi, Crit. Rev. Food Sci. Nutr., 2024, 64, 3139–3150.

[48] H. Carlsen, A.-M. Pajari, Food Nutr. Res., 2023, 67, 1–19.

[49] S. K. Gill, M. Rossi, B. Bajka, K. Whelan, Nat. Rev. Gastroenterol. Hepatol., 2021, 18, 101–116.

[50] F. De Vadder, P. Kovatcheva-Datchary, D. Goncalves, J. Vinera, C. Zitoun, A. Duchampt, F. Bäckhed, G. Mithieux, Cell, 2014, 156, 84–96.

[51] J. Vily-Petit, M. Soty-Roca, M. Silva, M. Raffin, A. Gautier-Stein, F. Rajas, G. Mithieux, Gut, 2020, 69, 2193–2202.

[52] R. Aoki, M. Onuki, K. Hattori, M. Ito, T. Yamada, K. Kamikado, Y.-G. Kim, N. Nakamoto, I. Kimura, J. M. Clarke, T. Kanai, K. Hase, Microbiome, 2021, 9, 188.

[53] F. B. Morel, Q. Dai, J. Ni, D. Thomas, P. Parnet, P. Fança-Berthon, J. Nutr., 2015, 145, 2052–9.

[54] S. M. Shallangwa, A. W. Ross, P. J. Morgan, Front. Microbiol., 2025, 16, 1–16.

[55] K. Weitkunat, S. Schumann, K. J. Petzke, M. Blaut, G. Loh, S. Klaus, J. Nutr. Biochem., 2015, 26, 929–937.

[56] R. St-Amand, É. T. Ngo Sock, S. Quinn, J. M. Lavoie, D. H. St-Pierre, Lipids Health Dis., 2020, 19, 1–12.

[57] B. O. Schneeman, J. Nutr., 1999, 129, 1424S–7S.

[58] M. L. Fernandez, T. A. Wilson, K. Conde, M. Vergara-Jimenez, R. J. Nicolosi, J. Nutr., 1999, 129, 1323–32.

[59] L. A. Simons, S. Gayst, S. Balasubramaniam, J. Ruys, Atherosclerosis, 1982, 45, 101–8.

[60] M. A. Levrat-Verny, S. Behr, V. Mustad, C. Rémésy, C. Demigné, J. Nutr., 2000, 130, 243–8.

[61] J. M. Gee, N. A. Blackburn, I. T. Johnson, Br. J. Nutr., 1983, 50, 215–24.

[62] H. Andersson, Eur. J. Clin. Nutr., 1992, 46 Suppl 2, S69–76.

[63] M. Minekus, M. Jelier, J.-Z. Xiao, S. Kondo, K. Iwatsuki, S. Kokubo, M. Bos, B. Dunnewind, R. Havenaar, Biosci. Biotechnol. Biochem., 2005, 69, 932–8.

[64] J. Yang, H. Wei, Y. Lin, E. S. H. Chu, Y. Zhou, H. Gou, S. Guo, H. C. H. Lau, A. H. K. Cheung, H. Chen, K. F. To, J. J. Y. Sung, Y. Wang, J. Yu, Gastroenterology, 2024, 166, 323–337.e7.

[65] S. B. Singh, A. Carroll-Portillo, H. C. Lin, Microorganisms, 2023, 11, 1772.

[66] Q. Zeng, D. Li, Y. He, Y. Li, Z. Yang, X. Zhao, Y. Liu, Y. Wang, J. Sun, X. Feng, F. Wang, J. Chen, Y. Zheng, Y. Yang, X. Sun, X. Xu, D. Wang, T. Kenney, Y. Jiang, H. Gu, Y. Li, K. Zhou, S. Li, W. Dai, Sci. Rep., 2019, 9, 1–10.

[67] M. Arifuzzaman, T. H. Won, T. T. Li, H. Yano, S. Digumarthi, A. F. Heras, W. Zhang, C. N. Parkhurst, S. Kashyap, W. B. Jin, G. G. Putzel, A. M. Tsou, C. Chu, Q. Wei, A. Grier, R. Longman, G. Sonnenberg, E. Scherl, R. Sockolow, D. Lukin, R. Battat, T. Ciecierega, A. Solomon, E. Barfield, K. Chien, J. Ferreira, J. Williams, S. Khan, P. S. Chong, S. Mozumder, L. Chou, W. Zhou, A. Ahmed, C. Zhong, A. Joseph, J. Gladstone, S. Jensen, S. Worgall, C. J. Guo, F. C. Schroeder, D. Artis, Nature, 2022, 611, 578–584.

[68] E. Catry, L. B. Bindels, A. Tailleux, S. Lestavel, A. M. Neyrinck, J.-F. Goossens, I. Lobysheva, H. Plovier, A. Essaghir, J.-B. Demoulin, C. Bouzin, B. D. Pachikian, P. D. Cani, B. Staels, C. Dessy, N. M. Delzenne, Gut, 2018, 67, 271–283.

[69] S. Huang, S. Dong, L. Lin, Q. Ma, M. Xu, L. Ni, Q. Fan, Front. Pharmacol., 2023, 14, 1226448.

[70] J. Yang, S. Zhang, S. M. Henning, R. Lee, M. Hsu, E. Grojean, R. Pisegna, A. Ly, D. Heber, Z. Li, J. Nutr. Biochem., 2018, 52, 62–69.

[71] J. Hu, P. Guo, R. Mao, Z. Ren, J. Wen, Q. Yang, T. Yan, J. Yu, T. Zhang, Y. Liu, Diabetes. Metab. Syndr. Obes., 2022, 15, 3933–3947.

[72] C. Visuthranukul, S. Sriswasdi, S. Tepaamorndech, S. Chamni, A. Leelahavanichkul, Y. Joyjinda, V. Aksornkitti, S. Chomtho, Int. J. Obes. (Lond)., 2024, 48, 1696–1704.

[73] J. Pessoa, G. D. Belew, C. Barroso, C. Egas, J. G. Jones, Nutrients, 2023, 15.

[74] J.-Z. Ye, Y.-T. Li, W.-R. Wu, D. Shi, D.-Q. Fang, L.-Y. Yang, X.-Y. Bian, J.-J. Wu, Q. Wang, X.-W. Jiang, C.-G. Peng, W.-C. Ye, P.-C. Xia, L.-J. Li, World J. Gastroenterol., 2018, 24, 2468–2481.

[75] A. H. Lichtenstein, Ann. Med., 1990, 22, 49–52.

[76] C. Juste, P. Gérard, Microorganisms, 2021, 9.

[77] L. Li, K. K. Buhman, P. A. Hartman, D. C. Beitz, Lett. Appl. Microbiol., 1995, 20, 137–40.

[78] L. Li, S. M. Batt, M. Wannemuehler, A. Dispirito, D. C. Beitz, Lab. Anim. Sci., 1998, 48, 253–5.

[79] J. G. M. Markle, D. N. Frank, S. Mortin-Toth, C. E. Robertson, L. M. Feazel, U. Rolle-Kampczyk, M. von Bergen, K. D. McCoy, A. J. Macpherson, J. S. Danska, Science, 2013, 339, 1084–8.

