## Supplementary Files for "Soluble and insoluble dietary fibres differentially affect liver steatosis and gut microbiota in western-diet fed mice"

### Additional files for:

| | Chow diet CD cellulose | | Western-diet WD cellulose | | Chow diet CD inulin | | Western-diet WD inulin | |
|---|---|---|---|---|---|---|---|---|
| | gm% | kcal% | gm% | kcal% | gm% | kcal% | gm% | kcal% |
| Protein | 20.8 | 23.44 | 42.75 | 39.38 | 20.8 | 22.79 | 20.8 | 18.73 |
| Carbohydrate | 58.95 | 66.42 | 42.75 | 39.38 | 65.7 | 67.35 | 49.5 | 40.76 |
| Fat | 4.00 | 10.14 | 20.00 | 41.46 | 4 | 9.86 | 20 | 40.51 |
| Total kcal | | 100 | | 100 | | 100 | | 100 |
| kcal/gm | 3.55 | | 4.34 | | 3.65 | | 4.44 | |
| **Ingredient** | **gm** | **kcal** | **gm** | **kcal** | **gm** | **kcal** | **gm** | **kcal** |
| Casein | 20.00 | 80.0 | 20.00 | 80.00 | 20.00 | 80.00 | 20.00 | 80.00 |
| Methionine | 0.30 | 1.20 | 0.30 | 1.20 | 0.30 | 1.20 | 0.30 | 1.20 |
| Corn starch | 35.95 | 143.80 | - | - | 35.95 | 143.80 | - | - |
| Maltodextrin | 15.50 | 62.00 | 11.75 | 47.00 | 15.5 | 62.00 | 11.75 | 47.00 |
| Sucrose | 7.00 | 28.00 | 30.50 | 122.00 | 7.00 | 28.00 | 30.50 | 122.00 |
| Cellulose | 9.00 | - | 9.00 | - | 2.25 | - | 2.25 | - |
| Inulin (Orafti® IPS) | - | - | - | - | 6.75 | 10.13 | 6.75 | 10.13 |
| Soybean oil | 4.00 | 36.00 | - | - | 4.00 | 36.00 | - | - |
| (13% vegetable oil + 7% olive oil) | - | - | 20.00 | 180.00 | - | - | 20.00 | 180.00 |
| Cholesterol | - | - | 0.2 | - | - | - | 0.20 | |
| Mineral PM 205B 7% Safe | 7.00 | - | 7.00 | - | 7.00 | - | 7.00 | - |
| Vitamin PV 200 1% Safe | 1.00 | 4.00 | 1.00 | 4.00 | 1.00 | 4.00 | 1.00 | 4.00 |
| Choline | 0.25 | - | 0.25 | - | 0.25 | - | 0.25 | - |
| **Total** | **100.00** | **355.00** | **100.00** | **434.20** | **100.00** | **365.13** | **100.00** | **444.33** |

**Table S1:** Composition of semi-synthetic experimental chow diet (CD) or western-diet (WD) supplemented with 9% dietary fibre.

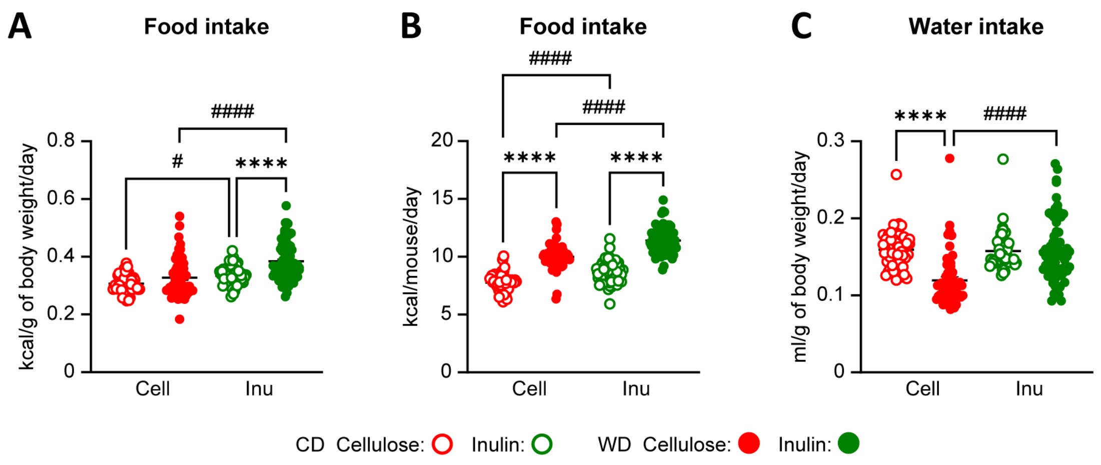

**Figure S1: Food and water consumption.**

*Food (A) and water (B) intake. Data are presented as the mean ± SEM. * diet effect or # fiber effect. *P < 0,05, ##P < 0,01, **** or ####P < 0,0001. Differential effects were analysed by analysis of variance (one-way ANOVA) with post hoc Šídák's test.*

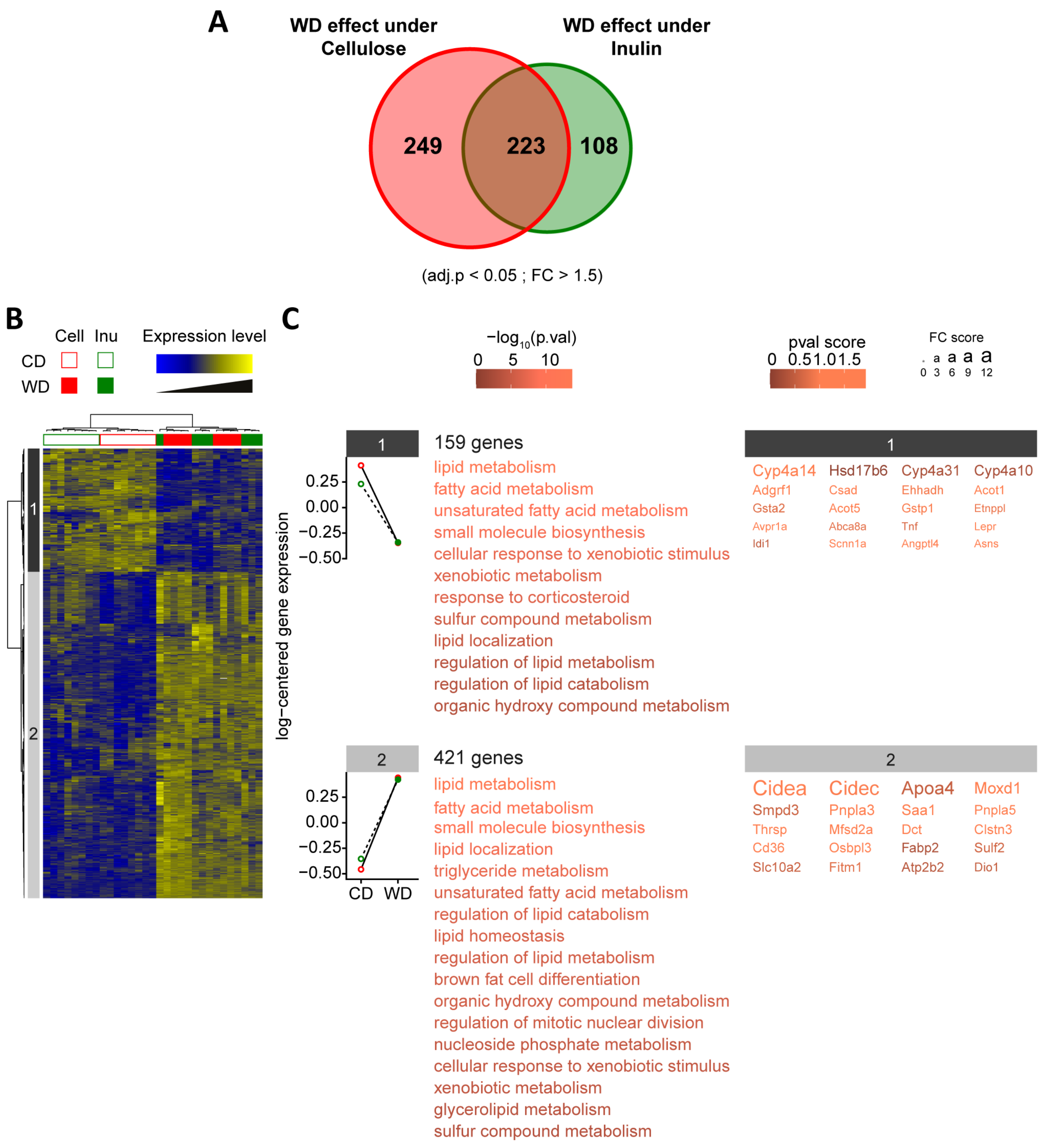

**Figure S2: Hepatic gene expression.**

*(A) Venn diagram representing the number of genes significantly regulated by diet (WD vs. CD) within each fibre (cellulose or inulin). (B) Heatmap of the 580 DEGs with a significant effect. (C) Mean expression profiles for the two gene clusters. The most significantly enriched biological processes and the 20 genes showing the largest differences in expression in each cluster are shown at the right of each profile.*

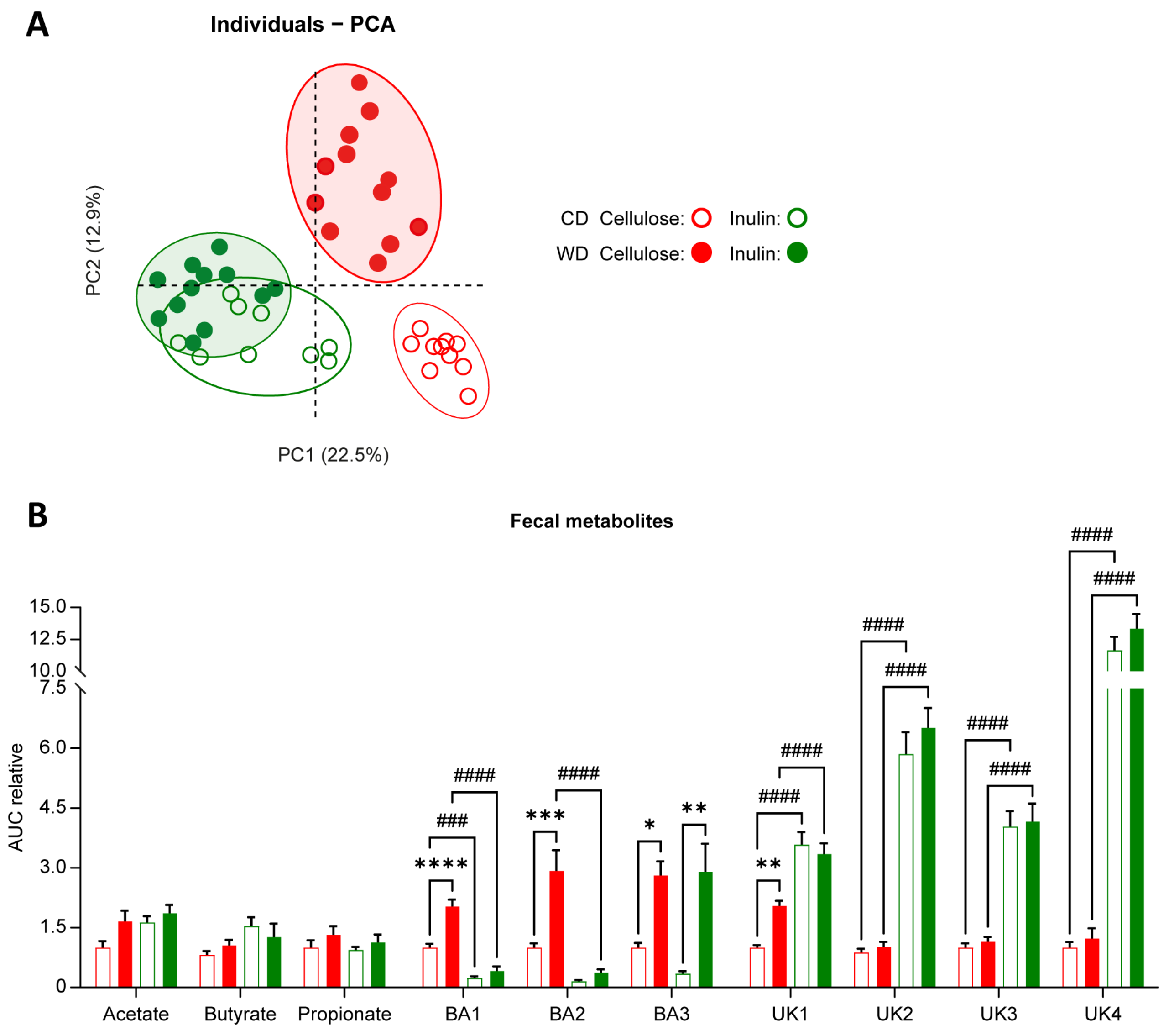

**Figure S3: Fecal metabolic profiles.**

*(A) Principal-component analysis (PCA) score plots of the whole-fecal metabolome datasets (n= 12/group). Each dot represents an observation (animal), projected onto first (horizontal axis) and second (vertical axis) PCA variables. (B) AUC of the $^{1}$-NMR spectra was integrated for short chain fatty acids (acetate, butyrate, propionate), bile acids (BA) and unknown aromatic metabolites (UK)*

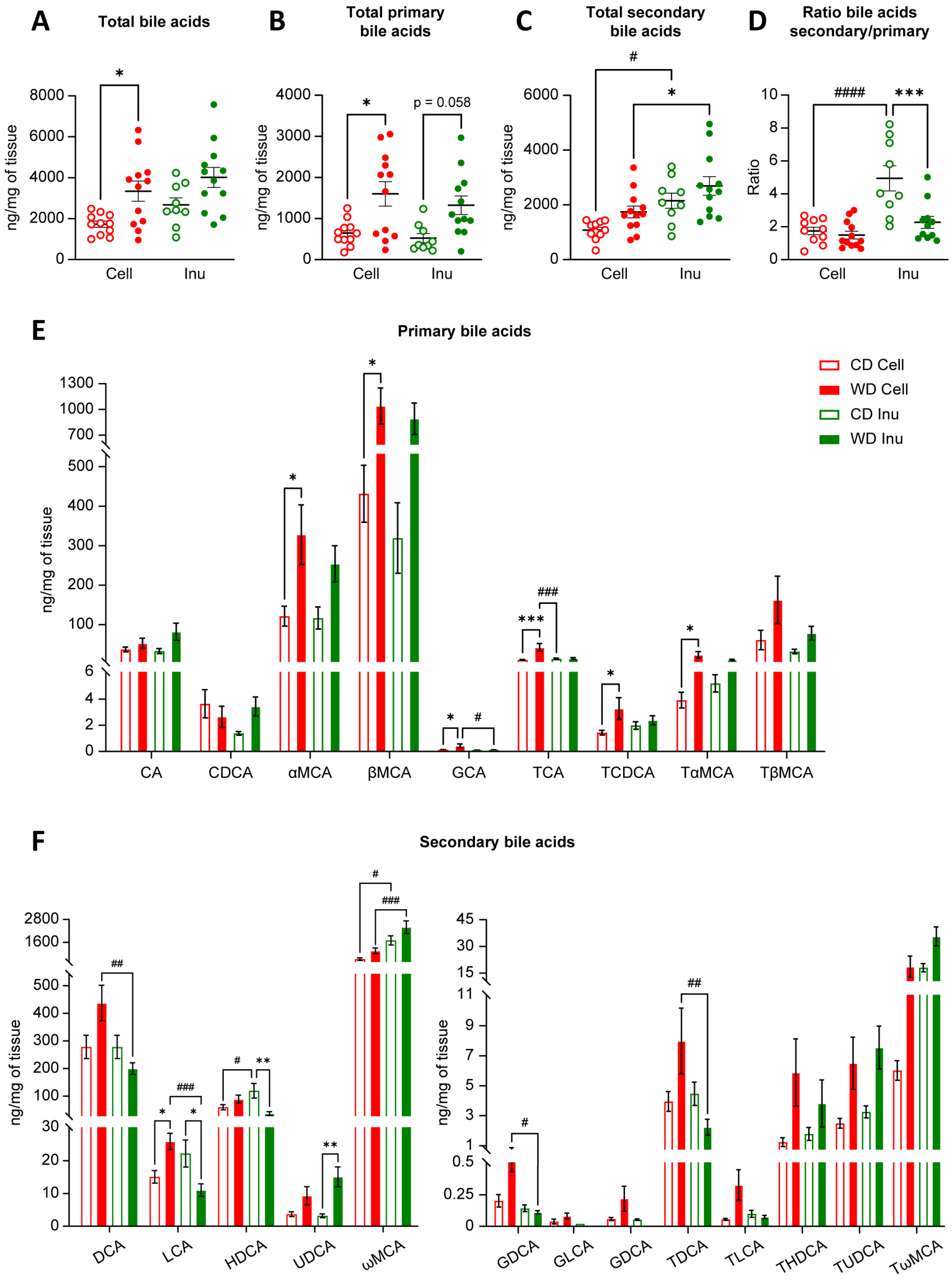

**Figure S4: Fecal bile acids.**

*(A-C) Total bile acids (B) total primary bile acids (C) total secondary bile acids concentrations extracted from feces and analysed by LC-MS. (D) Ratio secondary/primary bile acids (E) Individual primary bile acids (F) Individual secondary bile acids. Data are presented as the mean ± SEM for n = 9-12/group. * diet effect or # fiber effect. * or #P < 0,05, ** or ##P < 0,01, *** or ###P < 0,001, **** or ####P < 0,0001.*

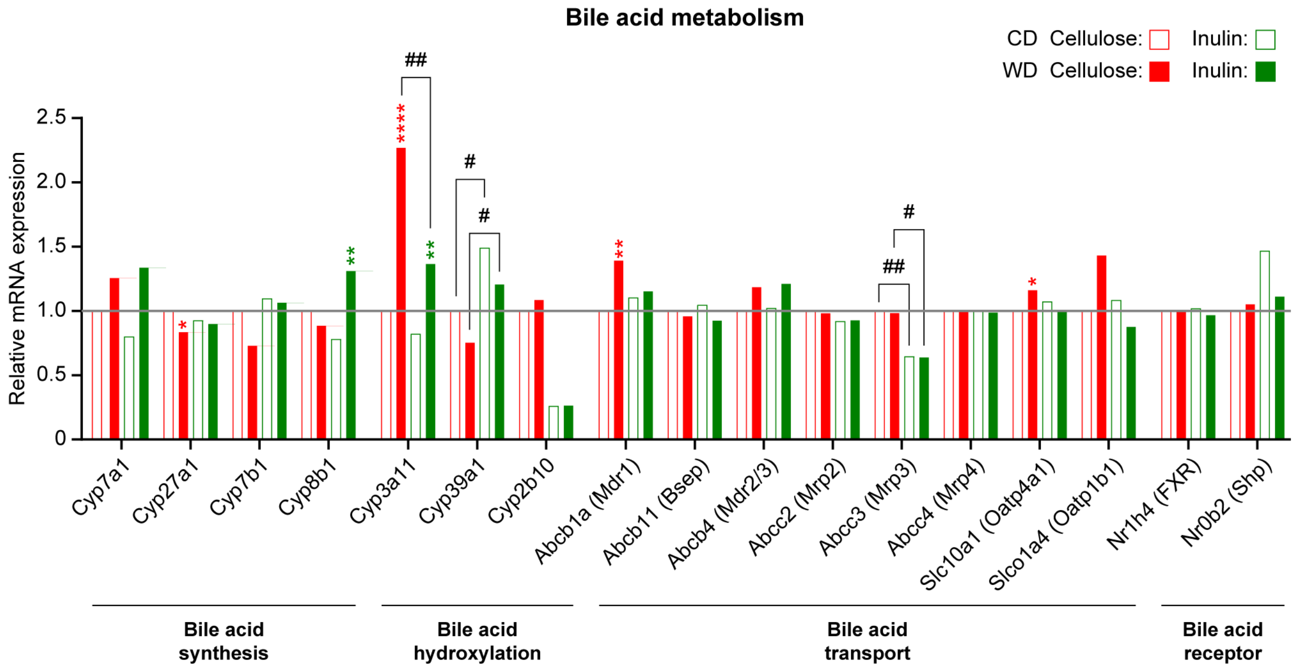

**Figure S5: Hepatic mRNA expression of genes involved in bile acid metabolism.**

*Hepatic mRNA expression involved in bile acid metabolism measured by microarray. Data are presented as the mean for n = 7-8/group. * diet effect or # fibre effect. * or #P < 0,05, ** or ##P < 0,01, *** or ###P < 0,001, **** or ####P < 0,0001.*
